# Neuronal identity control at the resolution of a single transcription factor isoform

**DOI:** 10.1101/2024.06.14.598883

**Authors:** Natalie Smolin, Mark Dombrovski, Bryce W. Hina, Anthony Moreno-Sanchez, Ryan Gossart, Catherine R. Carmona, Aadil Rehan, Roni H. Hussein, Parmis Mirshahidi, Jessica Ausborn, Yerbol Z. Kurmangaliyev, Catherine R. von Reyn

**Author notes:** Equal Contribution (Mark Dombrovski, Bryce W. Hina).

## Abstract

The brain exhibits remarkable neuronal diversity which is critical for its functional integrity. From the sheer number of cell types emerging from extensive transcriptional, morphological, and connectome datasets, the question arises of how the brain is capable of generating so many unique identities. ‘Terminal selectors’ are transcription factors hypothesized to determine the final identity characteristics in post-mitotic cells. Which transcription factors function as terminal selectors and the level of control they exert over different terminal characteristics are not well defined. Here, we establish a novel role for the transcription factor *broad* as a terminal selector in *Drosophila melanogaster*. We capitalize on existing large sequencing and connectomics datasets and employ a comprehensive characterization of terminal characteristics including Perturb-seq and whole-cell electrophysiology. We find a single isoform *broad-z4* serves as the switch between the identity of two visual projection neurons LPLC1 and LPLC2. *Broad-z4* is natively expressed in LPLC1, and is capable of transforming the transcriptome, morphology, and functional connectivity of LPLC2 cells into LPLC1 cells when perturbed. Our comprehensive work establishes a single isoform as the smallest unit underlying an identity switch, which may serve as a conserved strategy replicated across developmental programs.

## INTRODUCTION

The determination of a neuron’s identity—including the morphology, connectivity, and gene expression patterns that differentiate a neuronal cell—is a complicated task. Even within the humble fruit fly *Drosophila melanogaster*, more than 4,000 neural cell types (Scheffer et al., 2020; Schlegel et al., 2023) need to be determined, ensuring they exist in the appropriate space within the brain, establish their correct morphology, and connect to their intended partners, all within the volume of 5 nL (Makos et al., 2009). Our understanding of the regulatory logic required to specify unique and final neuronal identities is currently limited.

Conserved across vertebrates and invertebrates alike, neural progenitors produce cell types based on their location and age. Both spatially restricted transcription factors and temporal cascades of transcription factors cooperate to determine which cell types are generated at what particular time during development (reviewed in (Chen & Konstantinides, 2022; Holguera & Desplan, 2018; Kumar, 2001). Within the medulla of *Drosophila melanogaster*, for example, a stereotyped cascade of specific transcription factors (Hth➜Ey➜Slp➜D➜Tll) pattern developing neuroblasts, and loss of Slp expression in the sequence can impede the generation of entire populations of cell types (Li et al., 2013).

It is hypothesized that, in addition to transient transcription factor expression in progenitors, it is the persistent expression of certain transcription factors around the time of final mitosis throughout adulthood that specifies the terminal characteristics of a neuron (Allan & Thor, 2015; Hobert, 2008; Hobert, 2016; Holmberg & Perlmann, 2012). These ‘terminal selectors’ have been best studied in *C. elegans,* where selective removal or addition of transcription factors substantially altered the terminal characteristics, such as morphology and neurotransmitter type, that define neuron identity (Flames & Hobert, 2009; O’Meara et al., 2010; Serrano-Saiz et al., 2013). Although evidence from *C. elegans* suggests a single terminal selector may be sufficient to define specific characteristics of identity, terminal selectors in larger animals are thought to act together in a combinatorial code (Allan & Thor, 2015; Arber et al., 1999; Hobert, 2016; Hobert & Kratsios, 2019; Tsuchida et al., 1994; Wolfram et al., 2014). To date, knowledge about which transcription factors serve as terminal selectors remains limited. Additionally, the extent to which one terminal selector affects multiple characteristics that define type identity has not been well explored. Terminal selector studies are often limited to quantifying one or two characteristics to assume an identity change, even though characteristics are not necessarily linked (Konstantinides et al., 2018). For example, although perturbations of individual terminal selectors can substantially alter cellular morphology, changes in morphology may not affect connectivity, as neuronal partners can adapt to find intended partners even when morphology is atypical (Valdes-Aleman et al., 2021). More comprehensive evaluations of terminal selectors are therefore needed to understand the extent to which terminal selectors exert their control.

The *Drosophila* brain provides an excellent model for the study of terminal neuronal fate specification, due to its well-characterized, stereotyped wiring patterns and experimental tractability (Dorkenwald et al., 2022; Fischbach & Dittrich, 1989; Nern et al., 2015; Scheffer et al., 2020; Takemura et al., 2013; Takemura et al., 2015; Zheng et al., 2018). Electron microscopy datasets have enabled researchers to define the connectivity of a specific cell type (Dorkenwald et al., 2022; Nern et al., 2024; Scheffer et al., 2020; Schlegel et al., 2023; Zheng et al., 2018). Sophisticated genetic tools enable a ‘plug-and-play’ framework for inducing specific genetic perturbations exclusively in designated cell types (Dionne et al., 2018; Jenett et al., 2012; Lai & Lee, 2006; Perkins et al., 2015; Pfeiffer et al., 2010). *Drosophila* neurons are also amenable to whole-cell electrophysiology, enabling the functional consequences of perturbations to be evaluated (von Reyn et al., 2014; von Reyn et al., 2017). And perhaps most critically, recent advancements within the field of transcriptomics allow tracking of genetic expression in a cell-type-specific manner over the course of development, elucidating the temporal expression profile of specific genes (Davis et al., 2020; Konstantinides et al., 2018; Kurmangaliyev et al., 2020; Ozel et al., 2022; Ozel et al., 2021).

One recent single-cell RNA sequencing (scRNA-seq) dataset of the developing fly optic lobes (Kurmangaliyev et al., 2020) has uncovered sustained differential ‘on-off’ expression of the transcription factor *broad* in two visual projection neuron cell types (VPNs) called lobula plate lobula columnar neuron type 1 (LPLC1) and lobula plate lobula columnar neuron type 2 (LPLC2) (Figure 1). VPNs detect specific visual features, with LPLC1 and LPLC2 encoding features of an object approaching on a direct collision course (Ache et al., 2019; Klapoetke et al., 2017; Klapoetke et al., 2022; Tanaka & Clark, 2022; von Reyn et al., 2017). LPLC1 and LPLC2 innervate the same brain regions, extending dendrites within the lobula and lobula plate of the optic lobes, and projecting axons out of the optic lobes and into the ventral lateral protocerebrum (VLP) in the central brain. Within these brain regions, LPLC1 and LPLC2 differ in dendrite and axonal targeting (Figure 1a,b). LPLC1 dendrites target layers 2-5B of the lobula (Lo2-5B) and layers 2-4 of the lobula plate (LoP2-4), while LPLC2 dendrites target Lo4-5B and Lop1-4. LPLC1 and LPLC2 axons terminate in their own, separate glomeruli within the VLP. The resolvable differences in morphology along with the genetic accessibility of LPLC1 and LPLC2 make these cell types promising for investigating *broad* as a potential terminal selector that may differentiate the two cell types.

**Figure 1:**
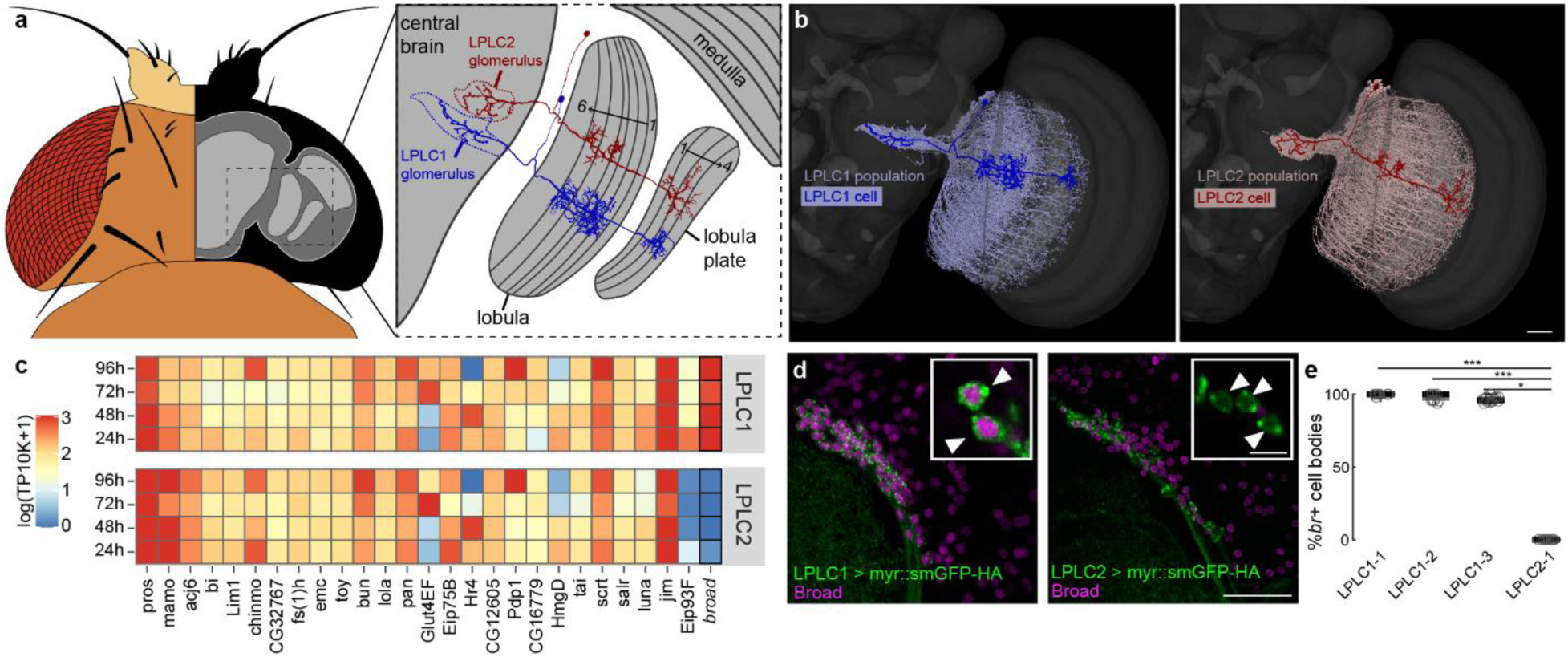
The transcription factor *broad* is differentially expressed between LPLC1 and LPLC2. **(a)** Fly brain schematic dorsal view illustrating single LPLC1 and LPLC2 neuron projection patterns. **(b)** Mesh reconstruction (Dorkenwald et al., 2022) of LPLC1 (left) and LPLC2 (right) populations within an anterior view of the fly brain. A single LPLC1 (left) and LPLC2 (right) drawing has been overlaid onto these populations. **(c)** Heatmaps showing expression of transcription factors in developing LPLC1 and LPLC2 neurons, data from (Kurmangaliyev et al., 2020). Columns are highly expressed genes that define these two cell-types, and rows indicate hours after pupae formation (APF). The average normalized expression is shown as log-transformed TP10K values (transcripts-per-10,000 UMI). See Methods for details. **(d)** Broad protein labeling displaying *broad* positive LPLC1 cell bodies (left) and *broad* negative LPLC2 cell bodies (right). Scale bar = 20 µm. Insets feature a zoomed in single plane with arrowheads pointing to individual cell bodies. Inset scale bar = 5 µm. **(e)** Quantification of *broad* positive somata in three LPLC1 driver lines and one LPLC2 driver line. N ≥ 6 animals for all conditions. Kruskal Wallis (p = 7.31e-8), Dunn-Sidak post hoc. * = p<0.05, ** = p<0.01, *** = p<0.001.

In *Drosophila*, the transcription factor *broad* has been well studied as having a key role during fly metamorphosis, and is necessary for the transition from larvae to pupae (Bayer et al., 1996; Karim et al., 1993; Restifo & White, 1991). *Broad* is expressed across tissues in the prepupal stages, induced through the global ecdysone hormone cascade that drives metamorphic events (Bayer et al., 1996; Brennan et al., 2001; Emery et al., 1994; Huet et al., 1993; Mugat et al., 2000; Zhou et al., 2009). *Broad* has also recently been implicated in the early cell patterning of type II neuroblasts (El-Danaf et al., 2023), working as a temporal transcription factor. Despite being well studied within the early stages of pupal development, very little is known of its ability to terminally differentiate cells.

Here, we investigate the role of the transcription factor *broad* as a terminal selector that differentiates two closely related VPNs during development, evaluating putative changes in cellular identity that result through perturbations of this program. Through the knock-down of *broad* in LPLC1 cells and the overexpression of *broad* in LPLC2 cells, we investigate morphological changes that indicate altered terminal identity. We establish differences between LPLC1 and LPLC2 synaptic partners using connectome datasets and use whole-cell patch clamp electrophysiology and optogenetics to determine how perturbing *broad* expression alters synaptic partner specificity and functional connectivity. Finally, we perform single-cell RNA sequencing after *broad* perturbation to evaluate the impact of individual *broad* isoforms on transcriptional identity. Our work elucidates mechanisms that define terminal cellular identity, establishing a role for *broad* as a terminal selector and revealing a minimal unit required to distinguish final and unique cell type identity.

## RESULTS

### *broad* is differentially expressed between LPLC1 and LPLC2

To identify candidate terminal selectors that may control differential terminal characteristics between LPLC1 and LPLC2, we referenced the scRNA-seq transcriptional atlas of the *D. melanogaster* visual system (Kurmangaliyev et al., 2020) and identified all transcription factors that are highly expressed in the developing LPLC1 and LPLC2 neurons (Figure 1c). Only two transcription factors, Eip93F and *broad*, were differentially expressed between these two cell types. However, only *broad* had an ‘on-off’ expression pattern that was maintained throughout development (Figure 1c), giving us reason to believe *broad* may help differentiate these two cell types. The time points 24-96 hours after pupal formation (h APF) are when LPLC1 and LPLC2 establish their morphology and wire into circuits (McFarland et al., 2024). We therefore hypothesized that *broad* is a key factor in differentiating LPLC1 from LPLC2, acting as a terminal selector to establish LPLC1 identity. We then validated that *broad* protein expression patterns matched the on-off mRNA expression patterns from scRNA-seq data with immunolabeling, confirming *broad* was only expressed in LPLC1 cell bodies (Figure 1d,e).

### Changes in *broad* expression induce morphological changes in LPLC1 and LPLC2

If *broad* is acting as a terminal selector to differentiate LPLC1 from LPLC2, perturbations in *broad* expression should directly impact the wiring programs of these cells, for instance, causing LPLC1 to resemble LPLC2. To investigate our terminal selector hypothesis, we knocked down *broad* in LPLC1 using two different RNAi lines that target all *broad* isoforms (Perkins et al., 2015).

We found both *broad* knock-down conditions led to alterations in LPLC1 axon morphology and targeting. The total volume occupied by axon terminals for the *broad* RNAi-1 condition increased slightly (Figure 2a,b), suggesting that these axons are occupying more space, extending outside of the LPLC1 glomerulus region. Surprisingly, we observed a decrease in total volume occupied by axon terminals for the *broad* RNAi-2 condition (Figure 2a,b), accompanied by an overall decrease in the number of LPLC1 somata (Supplementary Figure 1). We reasoned this decrease may occur because a portion of perturbed neurons are no longer fully being recognized by the LPLC1 driver line when *broad* expression is decreased. We found fluorescence within the LPLC1 glomerulus was not significantly changed after *broad* knock-down in LPLC1 (Figure 2c,d). Both knock-down conditions did however lead to LPLC1 axonal tracts targeting the LPLC2 glomerulus (Figure 2e,f). These data suggest *broad* expression in LPLC1 establishes its axon morphology, and the loss of *broad* in LPLC1 redirects its axons towards LPLC2 targets.

**Figure 2:**
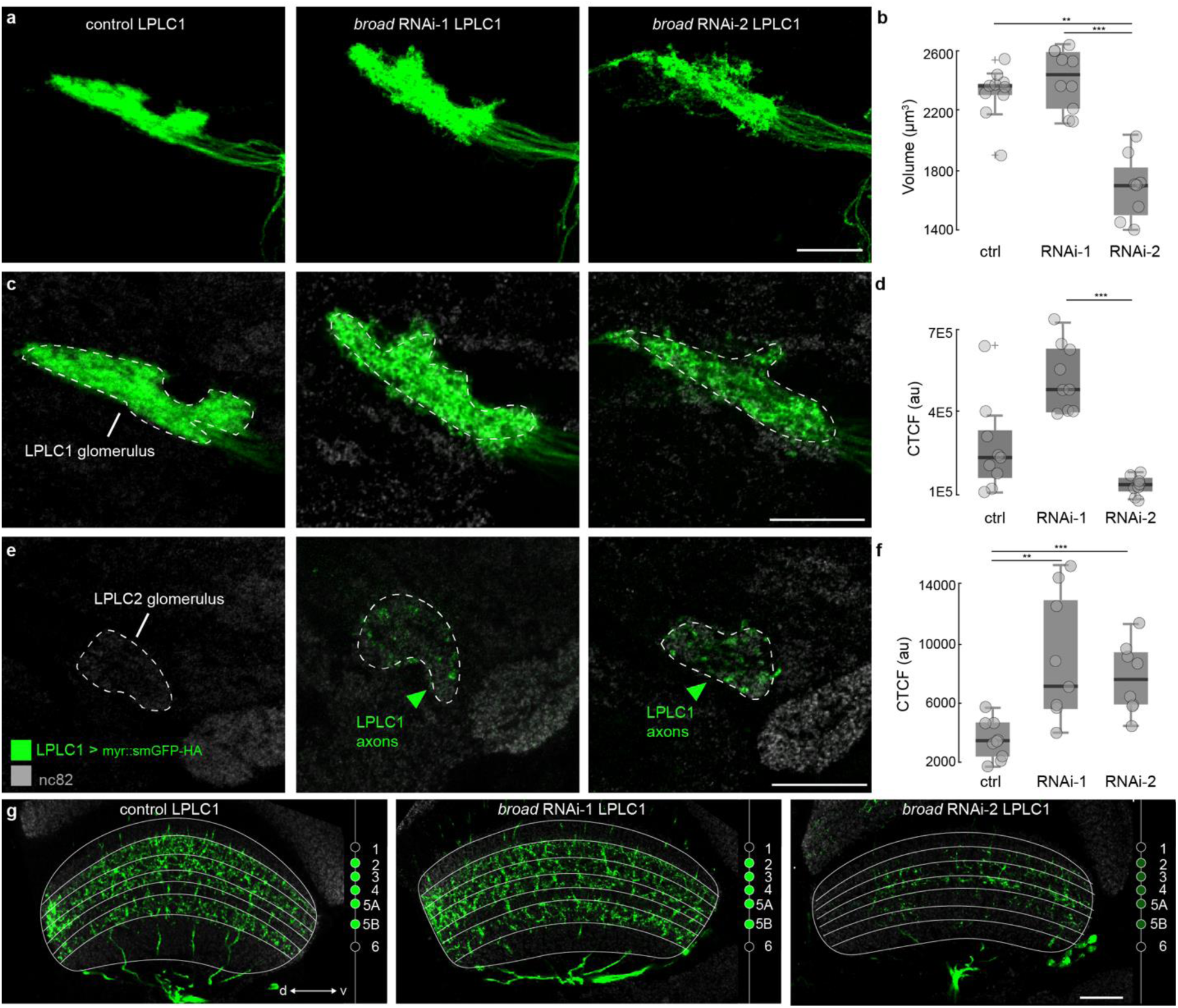
Knock-down of *broad* in LPLC1 cells results in ectopic axonal branching into the LPLC2 glomerulus. **(a)** Maximum projection images of axons tracts for control LPLC1 and *broad* knock-down (*broad* RNAi-1 and *broad* RNAi-2) LPLC1 cells. Scale bar = 20 µm. **(b)** Quantification of overall axonal volume of each condition in **(a)**. N ≥ 4 animals for each condition. Kruskal Wallis (p = 2.98e-04), Dunn-Sidak post hoc. * = p<0.05, ** = p<0.01, *** = p<0.001. **(c)** Single plane images depicting the innervation of the LPLC1 glomerulus by LPLC1 axons. Scale bar = 20 µm. **(d)** Quantification of LPLC1 axon density (corrected total cell fluorescence, CTCF) in the LPLC1 glomerulus shown in **(c)**. N ≥ 4 animals for each condition. Kruskal Wallis (p = 1.84e-04), Dunn-Sidak post hoc * = p<0.05, ** = p<0.01, *** = p<0.001. **(e)** Single plane images depicting the innervation of the LPLC2 glomerulus by LPLC1 axons. Scale bar = 20 µm. **(f)** Quantification of LPLC1 axon density (CTCF) in the LPLC2 glomerulus shown in **(e)**. N ≥ 4 animals for each condition. Kruskal Wallis (p = 0.0053), Dunn-Sidak post hoc * = p<0.05, ** = p<0.01, *** = p<0.001. Scale bar = 20 µm.

We note we did not witness a full targeting switch for the axons, which could be due to physical competition with LPLC2 for space on postsynaptic partners (McFarland et al., 2024). We also witnessed no morphological changes to LPLC1 dendrites with *broad* knock-down (Figure 2g), which could signify a limiting time-lag between RNAi expression and subsequent protein knock-down (Yao et al., 2015). Terminal selectors are thought to require early and sustained expression across development, and as our split-GAL4 LPLC1 line turns on at around 12h-24h APF (McFarland et al., 2024), these RNAi tools may not start working early enough to completely alter axon targeting and dendrite morphology.

Based on our finding that the knock-down of *broad* changes LPLC1 axonal projection targeting towards LPLC2 targets in the central brain, we next investigated whether the overexpression of *broad* could alter the morphology of LPLC2 to resemble LPLC1. We expressed individual isoforms of *broad* in LPLC2 and found only two (*broad-z3* and *broad-z4*) affected LPLC2 morphology (Figure 3 and Supplementary Figure 2). At the level of the dendrites, LPLC2 innervates layers 4-5B of the lobula, but *broad-z3* or *broad-z4* overexpression resulted in dendritic innervation across layers 2-5B of the lobula, which is the typical innervation pattern of LPLC1 (Figure 3a) (Wu et al., 2016).

**Figure 3:**
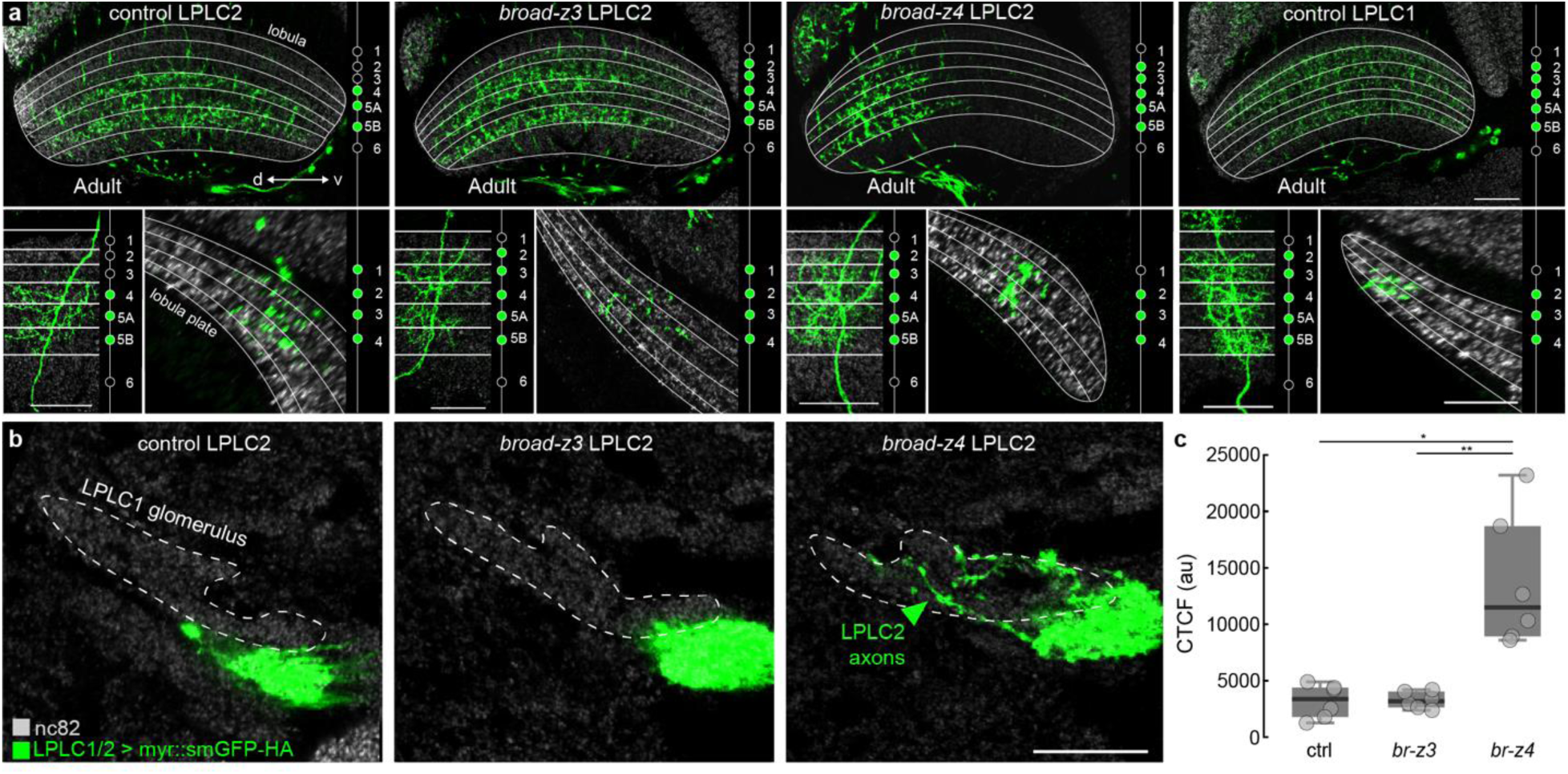
*broad-z3* and *broad-z4* overexpression in LPLC2 cells result in LPLC1-like morphologies. **(a)** (top) Innervation of the lobula for control LPLC2 cells, *broad-z3* LPLC2 cells, *broad-z4* LPLC2 cells, and control LPLC1 cells. Scale bar = 20 µm. (bottom left) Single cell morphology of cells within the lobula. Scale bar = 20 µm. (bottom right) Single cell morphology of cells within the lobula plate. N ≥ 6 animals for each condition. Scale bar = 20 µm. **(b)** Innervation of the LPLC1 glomerulus for control, *broad-z3*, and *broad-z4* LPLC2 cells. **(c)** Quantification of LPLC2 axon density (CTCF) in the LPLC1 glomerulus shown in **(b)**. N ≥ 6 animals for each condition. Kruskal Wallis (p = 0.0034), Dunn-Sidak post hoc * = p<0.05, ** = p<0.01, *** = p<0.001. Scale bar = 20 µm.

To better resolve individual morphologies, we used the MultiColor FlpOut (MCFO) technique (Nern et al., 2015) to sparsely label control LPLC1 cells, control LPLC2 cells, *broad-z3* LPLC2 cells and *broad-z4* LPLC2 cells (Figure 3a). As observed when labeling the full population, we found at the single cell level *broad-z3* and *broad-z4* overexpression in LPLC2 cells caused the lobula dendrites to exhibit LPLC1-like morphology. Sparse labeling also enabled us to resolve dendrite arborization patterns in the lobula plate. We found control LPLC2 cells to arborize in LoP1-4 and control LPLC1 cells to arborize in LoP2-4 (Figure 3a), consistent with current literature (Tanaka & Clark, 2022; Wu et al., 2016). Furthermore, we found that only *broad-z4* changed LPLC2 dendritic innervations pattern within the lobula plate to LoP2-4, resembling LPLC1 dendrites.

We next investigated how *broad-z3* and *broad-z4* overexpression changes the axonal morphology of LPLC2 cells. We found *broad-z4* LPLC2 axons innervated the LPLC1 glomerulus (Figure 3b,c), suggesting the expression of *broad-z4* is sufficient to redirect LPLC2 axons towards LPLC1 postsynaptic targets. Overall, *broad-z4* demonstrated the most significant alterations in terminal LPLC2 morphology across the lobula, lobula plate, and axonal terminals.

We also note that *broad-z4* overexpression in adult LPLC2 cells consistently appeared to lead to a loss of dendrites in the dorsal half of the lobula and lobula plate (Figure 3a), and a decrease in the overall number of LPLC2 somata. Upon further examination, however, we found the “missing” population was weakly labeled by the LPLC2 driver line, indicating the driver line may have difficulty recognizing these perturbed cells as still being LPLC2 (Supplementary Figure 3).

### LPLC2 cells have altered connectivity resulting from *broad* overexpression

The observed alterations in layer specific dendrite targeting suggest there may be a change in synaptic partners that occurs when *broad* is expressed in LPLC2, with LPLC2 receiving inputs from lobula or lobula plate cell types normally reserved for LPLC1. To establish which changes in synaptic partners could occur with *broad* expression, we used the Full Adult Fly Brain (FAFB) electron microscopy (EM) dataset (Dorkenwald et al., 2022; Zheng et al., 2018), to determine the presynaptic inputs for LPLC1 and LPLC2 (Figure 4a). While we found Tm5f, Tm20, and Tm4 (Supplementary Figure 5) to provide strong inputs to both LPLC1 and LPLC2 populations, we found T2 cells to provide one of the largest differences in lobula presynaptic inputs between LPLC1 and LPLC2 (Figure 4a,c). T2 neurons form synapses with LPLC1 in layers 2 and 3 of the lobula, but do not provide a major synaptic input to LPLC2. Since LPLC2 that overexpress *broad-z3* and *broad-z4* show novel dendritic innervations in layers 2 and 3 of the lobula, we hypothesized *broad* expressing LPLC2 may be forming direct, functional synapses with T2 in these layers.

**Figure 4:**
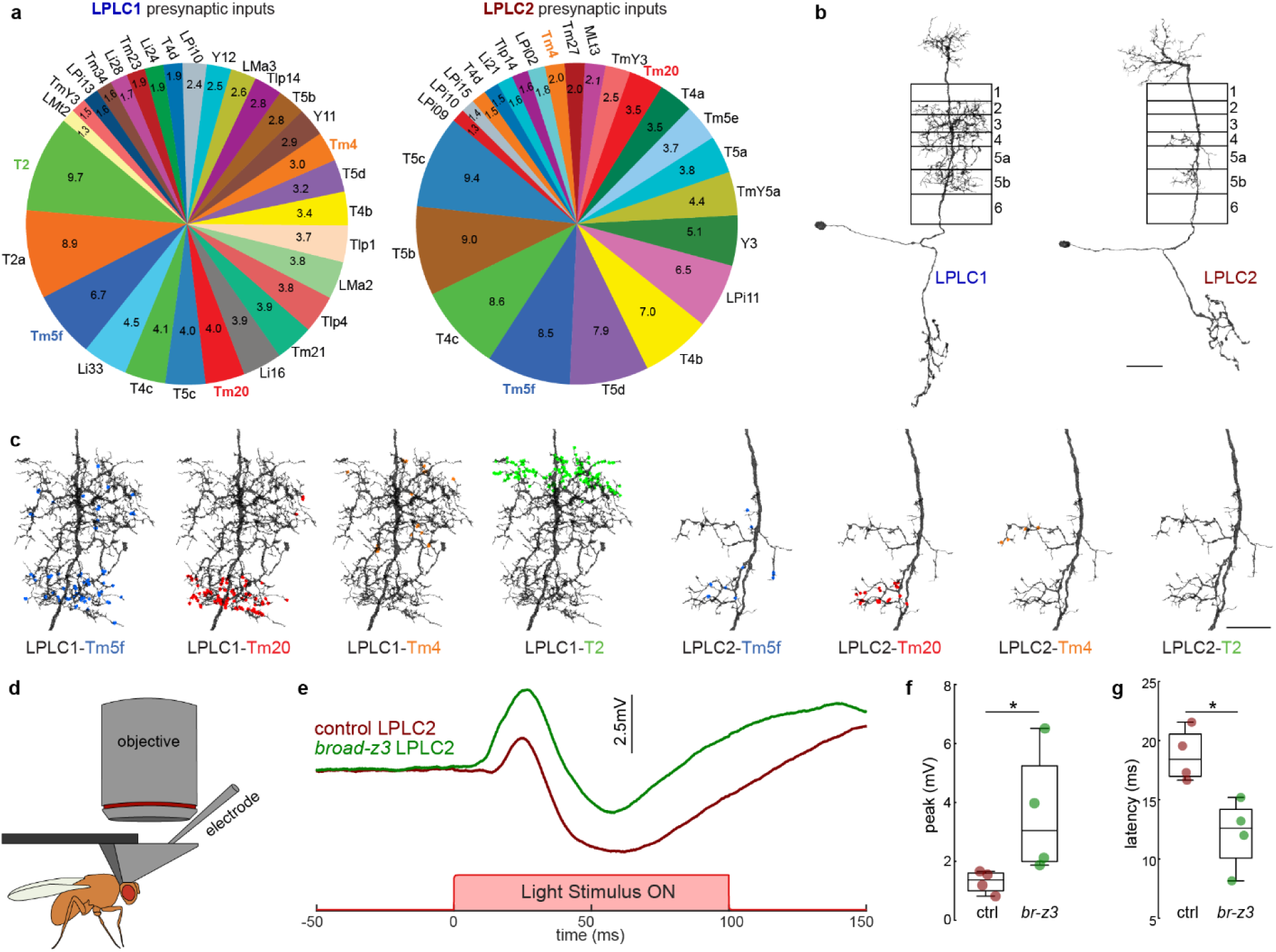
T2 cells are now connected to LPLC2 cells with the overexpression of *broad*. **(a)** Presynaptic inputs greater than 1% of total synapse counts to LPLC1 and LPLC2. **(b)** Mesh reconstruction (Dorkenwald et al., 2022; Schlegel et al., 2023; Zheng et al., 2018) of a representative (left) LPLC1 or (right) LPLC2 neuron. Approximate lobula layers have been boxed over the dendrites. Scalebar = 15 µm. **(c)** Mesh reconstructions of all synapses (colored circles) from Tm5f, Tm20, Tm4, and T2 neurons (Buhmann et al., 2021; Dorkenwald et al., 2022; Heinrich et al., 2018; Schlegel et al., 2023; Zheng et al., 2018) on a (left) LPLC1 and a (right) LPLC2 neuron. Scalebar = 10 µm. **(d)** Schematic illustrating whole-cell electrophysiology and optogenetics setup with light delivery through the objective. **(e)** Average responses for LPLC2 with and without *broad* isoform overexpression when T2 are optogenetically activated. **(f,g)** Quantification of **(f)** peak depolarization and **(g)** activation latency. N = 4 animals for each condition. Wilcoxon Rank Sum Test, * = p<0.05.

To investigate whether these putative synapses between T2 and *broad* expressing LPLC2 were functional, we combined optogenetic activation of T2 with whole-cell electrophysiology recordings from LPLC2 to assess direct synaptic coupling (Figure 4d,e). From these recordings, we found that LPLC2 cells expressing *broad-z3* have a significantly larger depolarization when T2 cells are optogenetically activated, as compared to control LPLC2 cells (Figure 4e,f). We also find *broad-z3* LPLC2 responses exhibit a shorter latency than control responses, suggesting direct synaptic coupling (Figure 4e,g). Our results suggest that the overexpression of *broad-z3* is sufficient to change connectivity in LPLC2 cells, and that LPLC2 cells are now functionally connected to T2 cells.

### *broad-z4* recodes transcriptomes of LPLC2 cells into LPLC1-like cells

Our data to this point suggest *broad* establishes aspects of morphology and functional connectivity of LPLC1 cells, distinguishing them from LPLC2, as predicted if *broad* is acting as a terminal selector. One outstanding question of terminal selectors is to what extent they determine identity. Do they only serve to control a subset of terminal characteristics, or do they have the capacity to fully switch cell type identity? We therefore investigated whether *broad*’s control extended beyond morphology to change the transcriptional profile of LPLC2 to resemble LPLC1. To investigate to what extent *broad* determines neuronal identity, we performed a single-cell RNA sequencing experiment of purified LPLC2 cells in wild-type (control) and *broad* overexpression conditions (Perturb-seq). To minimize technical variation between experimental conditions, we leveraged a genetic multiplexing strategy using wild-type chromosomes from the *Drosophila* Genetic Reference Panel (DGRP) strains as molecular tags for individual replicates (Figure 5a) (Huang et al., 2014; Kang et al., 2018; Mackay et al., 2012). This strategy was previously used for high-throughput multiplexed profiling of developmental trajectories of neurons (Kurmangaliyev et al., 2020) and genetic perturbations of neurons (Jain et al., 2022).

**Figure 5:**
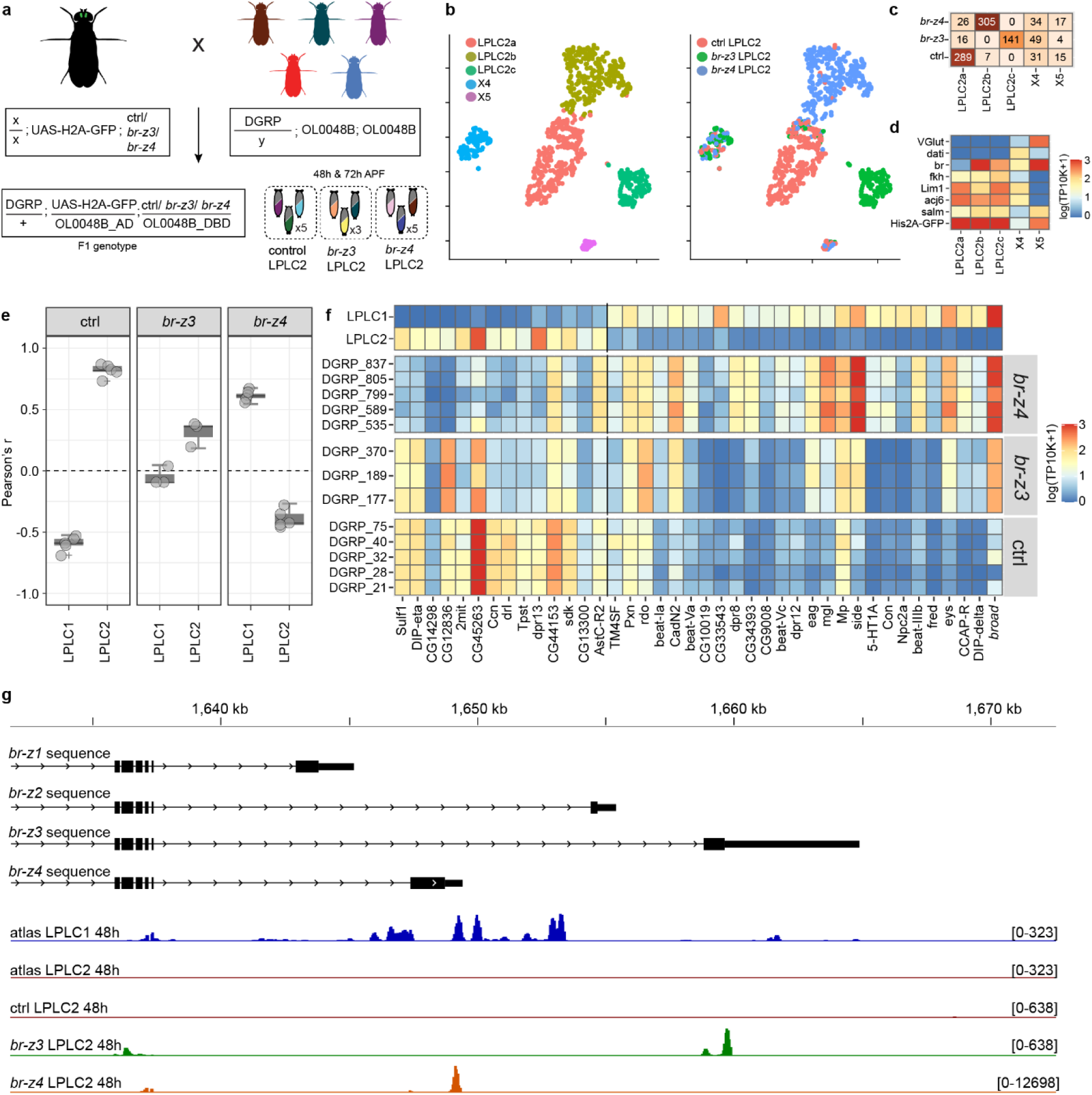
Overexpression of *broad-z4* recodes transcriptional identities of LPLC2 cells. **(a)** Experimental design of the multiplexed single-cell Perturb-Seq experiment. F1-generation of pupae carry the LPLC2 split-Gal4 driver (OL0048B), a nuclear GFP reporter (UAS-H2A-GFP), an overexpression or control construct (ctrl, *broad-z3*, *broad-z4*), and a replicate-specific wild-type X-chromosome (DGRP). LPLC2 neurons were purified and used for scRNA-seq at two timepoints (48h and 72h APF). Each experimental condition was replicated 3-5 times. The analysis at 48h APF is shown in **(b-g)**; the analysis at 72h APF is shown in Supplementary Figure 4. See Methods for more details. **(b)** t-distributed stochastic neighbor embedding (tSNE) plots are used only for visualization of the clustering of the data. Left, cells are color coded based on unsupervised clustering; Right, cells are color coded based on experimental conditions. **(c)** Cell counts across clusters and conditions. **(d)** Expression levels of LPLC2-specific transcription factors (Kurmangaliyev et al., 2020; Ozel et al., 2022) nuclear GFP reporter, and two genes enriched in ectopic, non-LPLC2, clusters (X4/X5). **(e)** Correlation analysis between transcriptional profiles of LPLC1 and LPLC2 neurons from the developmental atlas (Kurmangaliyev et al., 2020) and LPLC2 neurons in each experimental condition. Comparisons are based on differentially expressed genes (DEGs) between LPLC1 and LPLC2 from **(f)**. Circles are Pearson’s r for individual replicates; boxplots are distributions. **(f)** Heatmaps of expression patterns of DEGs between LPLC1 and LPLC2 at 48h APF. Expression patterns are shown in the atlas (top) and for each condition and replicate. DGRP lines for each X chromosome and replicate are indicated. See Methods for thresholds. **(g)** Coverage plots of scRNA-seq reads at the 3’-UTR region of *broad* in the atlas and Perturb-seq datasets. Most of the reads support the expression of the *broad-z4* isoform in LPLC1 neurons. Note that coverage of the overexpression constructs is restricted to the coding regions of corresponding transcripts.

We purified and sequenced LPLC2 cells overexpressing *broad-z3*, *broad-z4*, and a control genotype at 48h and 72h APF (Figure 5, Supplementary Figure 4), as many aspects of cell type identity and wiring patterns are established at these stages of development (Kurmangaliyev et al., 2020; McFarland et al., 2024). Overexpression of the two different *broad* isoforms resulted in two transcriptionally distinct populations of LPLC2 neurons indicating significant and distinct transcriptional effects of each isoform (Figure 5b-d, Supplementary Figure 4a-c).

Next, we compared expression profiles of *broad*-expressing LPLC2 neurons to wild-type LPLC1 and LPLC2 neurons obtained from the developmental atlas of the *Drosophila* visual system (Kurmangaliyev et al., 2020). The correlation analysis based on differentially expressed genes between LPLC1 and LPLC2 revealed distinct effects of the *broad* isoforms (Figure 5e-f). Transcriptomes of *broad-z3* LPLC2 neurons differed from both wild-type LPLC1 and LPLC2 neurons, but were more similar to wild-type LPLC2 neurons. In contrast, *broad-z4* LPLC2 neurons became more similar to wild-type LPLC1 than wild-type LPLC2 neurons. Taken together, our data suggest that *broad-z4* acts as a terminal selector between developing LPLC1 and LPLC2 cells, and acts as a key determining factor that establishes final transcriptional cell identity during these late-stages of neuronal development.

Given that our Perturb-seq data implicate *broad-z4* in switching identity between LPLC1 and LPLC2, we investigated what isoform is present natively in control LPLC1 cells. The *broad* gene is comprised of four isoforms, linking a common 5’ N-terminal to one of four pairs of 3’ zinc fingers. Since the atlas of the developing visual system generated 3’ RNA reads, we were able to determine which *broad* isoforms are expressed in each cell type. Four annotated isoforms of *broad* differ by zinc-finger domains encoded by alternative exons at 3’-UTR of the gene (Gramates et al., 2022). Excitingly, the analysis of scRNA-seq reads coverages in wild-type datasets showed that LPLC1 cells only express the *broad-z4* isoform (Figure 5g). These data corroborate our morphological studies, as *broad-z4* is the only isoform that induces morphological changes of LPLC2 at both their axons and dendrites. Our data suggest that while *broad* may be the distinguishing transcription factor between LPLC1 and LPLC2, the regulatory control it exerts is at the level of the isoform, and the differences between LPLC1 and LPLC2 cells are controlled by the *broad-z4* isoform.

## DISCUSSION

Terminal selector genes are thought to establish the terminal characteristics that define final cell identities (Hobert, 2008; Hobert, 2016). Final identity often requires a unique combinatorial expression of terminal selectors to implement all terminal characteristics (Allan & Thor, 2015; Hobert, 2016; Ozel et al., 2022). In this work, we elucidate one gene, *broad*, that is able to act as a terminal selector to differentiate neuronal identity between LPLC1 and LPLC2 cells through morphological, transcriptomic, and functional connectivity changes. Delving further, we find this control is exerted at the level of a single isoform, and that *broad-z4* is capable of fully switching the identity of LPLC2 cells.

We demonstrate a novel role for *broad*, working as a terminal selector to establish final and unique cellular identities within the fly visual system. *Broad* is best known to ensure appropriate stage-specific responses to hormone signaling during metamorphosis (Karim et al., 1993). Null mutations and deletions of *broad* lead to lethality at the end of the larval stage, indicating the requirement of *broad* for normal fly metamorphosis (Belyaeva et al., 1980; Kiss et al., 1988; Kiss et al., 1976; Restifo & White, 1991). More recent works have implicated *broad* as a temporal transcription factor, working in a cascade to pattern developing type II neuroblasts (El-Danaf et al., 2023). Through both temporal and spatial patterning, the nervous system is able to re-use the same genes in different contexts to allow a brain to overcome the wiring problem inherent to a brain—the idea that the number of connections in a brain outnumber the number of genes it has by several orders of magnitude (Hassan & Hiesinger, 2015). The function of *broad* as a terminal selector may be specific in converting LPLC2 into LPLC1 neurons, but *broad* may be capable of acting as a terminal selector across multiple cell types. Future work should investigate whether *broad* exerts control over the same terminal characteristics and utilizes the same mechanisms to exert its control.

We find *broad*’s terminal selector effects are specific to the individual *broad* isoform. *Broad* encodes four major protein isoforms, *broad-z1*, *-z2*, *-z3*, and *-z4*. These isoforms share a common animo-terminal core region but differ in their carboxy-terminal Zn fingers (Bayer et al., 1997; DiBello et al., 1991). *Broad*, formerly called *Broad*-Complex or BR-C, is known to encode for three functions throughout the *Drosophila* central nervous system: *br* (*broad*), *rbp* (reduced bristle number on palpus), and *2Bc* (Belyaeva et al., 1980; Kiss et al., 1988). All three functions control aspects of CNS reorganization: *br*^+^ is required for leg and wing imaginal disc morphogenesis and tanning and hardening of the larval cuticle, *rbp*^+^ is required for muscle and bristle development, *2Bc*^+^ is required for the closure of the thoracic epidermis (Bayer et al., 1997; Crossgrove et al., 1996). The *br* function is thought to be provided by the *-z2* isoform, the *rbp* function by the *-z1* and *-z4* isoforms, and the *2Bc* by the *-z3* isoform (Bayer et al., 1997). Here we define an additional, novel role for the *broad-z4* isoform as a terminal selector of LPLC1 cells. *Broad-z4* is the isoform present in native LPLC1 cells (Figure 5g) and perturbations of *broad-z4* result in the most significant switching of LPLC2 to LPLC1-like morphology at both the axons and the dendrites of these cells (Figure 3), as well as an LPLC1-like switch in transcriptional identity (Figure 5).

Interestingly, *broad-z3* also seems to have an effect on changing LPLC2 morphology and gene expression profiles, albeit to a lesser extent. Previous studies have indicated that there may be partial functional redundancy among the different *broad* isoforms. One type of *rbp* mutant (resulting in reduced bristle numbers) are fully rescued by the *-z1 broad* isoform but also partially rescued by the *-z4 broad* isoform (Bayer et al., 1997). Similarly, *2Bc* lethality can be rescued by *broad-z3* protein expression, but also partially rescued by *broad-z2* protein expression (Bayer et al., 1997). Here, we support these past findings that different *broad* isoforms have functional redundancies, but in a new context. In addition to the hydrophobic residues, cysteines, and histidines present in all C2H2 zinc-finger proteins (Miller et al., 1985), there is a high degree of similarity within the different zinc-finger domains (Bayer et al., 1996). The DNA binding domain of the *-z3* isoform, for instance, is quite similar to that of the *-z4* isoform (Bayer et al., 1996), and the redundancy that we see between the two isoforms could be based on structural similarity. *Broad-z4*, however, is the only isoform with a 5’ untranslated sequence of cDNA that diverges from the core sequence at position 163 kb (Bayer et al., 1996). This sequence specific to the *-z4* isoform may be part of the reason why the *-z4* isoform has such strong effects on changing terminal identity of LPLC2 cells. Previous studies have also found terminal selectors to act at the level of the isoform, however these studies have often focused their investigations on only a few aspects of terminal identity such as morphology, neurotransmitter identity, and the altered expression of downstream genes (Ahn et al., 2022; Campbell & Walthall, 2016; Chu et al., 2024; Neville et al., 2014; Pereira et al., 2019; Remesal et al., 2020).

Here, we quantitatively demonstrate how a terminal selector results in functional changes in synaptic specificity, expanding beyond changes in morphology and gene expression that have been a common read out for terminal selectors. Overexpression of an LPLC1 terminal selector, *broad*, in LPLC2 cells causes them to be synaptically coupled to T2 cells, which are LPLC1-specific synaptic partners. We anticipate *broad*-LPLC2 cells would have different visual tuning, as they now receive novel visual information, and would demonstrate responses matched to LPLC1 tuning for visual stimuli such as small moving squares or back-to-front motion (Klapoetke et al., 2022; Tanaka & Clark, 2022).

Terminal selectors are thought to activate cell-type specific effector genes that define the final differentiated identity of a mature neuron (Hobert & Kratsios, 2019). Differentially expressed genes from our Perturb-seq experiment may give insights as to target genes where *broad* exerts its regulatory control. Loss of *broad* results in altered expression of numerous genes, including cellular adhesion molecules that control synaptic specificity, as seen in our own Perturb-seq dataset. Other studies have found loss of a terminal selector to result in changes in gene expression including TFs, GPCRs, and ion channels (Hobert, 2016; Ozel et al., 2022; Wyler et al., 2016). Further chromatin binding studies such as ChiP-seq or DamID-seq would be required to definitively determine what downstream targets *broad* regulates.

Work *in C. elegans* has proposed terminal selectors for 98/118 neuron classes (Hobert, 2021), yet in other invertebrates and vertebrates, terminal selectors have bene identified in very few cell types. Our identification here of a terminal selector that specifies unique and final cell identities provides the most in-depth characterization of changes in identity that result from perturbing a terminal selector. The methodology here may also serve as a general framework as to how terminal selectors may be identified and evaluated in the future. The identification of terminal selectors enables further investigations into the mechanisms, downstream of terminal selectors, by which final cell identity characteristics are established. A failure to appropriately establish identity and maintain terminal characteristics may underlie neurodevelopmental and neuropsychiatric disorders, and many terminal selectors have orthologs in humans that have been involved in these disease states (Chao et al., 2017; Deneris & Hobert, 2014; Sahay & Hen, 2007; Sleven et al., 2017). In fact, the human ortholog of *broad* is BTBD18, and the genetic loci where it resides has recently been discovered to be linked to schizophrenia (Schizophrenia Working Group of the Psychiatric Genomics, 2014). The future impact of this work may therefore yield strategies to resolve or redefine neuronal identity in a clinical setting.

## MATERIALS AND METHODS

### Fly Stocks and developmental staging

*Drosophila melanogaster* were reared on a standard molasses, cornmeal, and yeast diet (Archon Scientific) and kept at 25°C and 60% humidity on a 12-hour light/dark cycle throughout development. For optogenetics experiments, larval flies were raised in the dark on low retinal food (standard food plus 0.2 mM retinal) and switched to high retinal food (standard food plus 0.4 mM retinal) upon eclosion. For developing pupal experiments, white pre-pupae (0h APF) were collected and incubated for the indicated number of hours. All experiments were performed on staged pupae or adult female flies 2-5 days post-eclosion. Fly genotypes are listed in Supplemental Table 1.

### Visualization of single-cell morphology

To visualize single cells of LPLC1 and LPLC2, we used MultiColor FlpOut (MCFO) (Nern et al., 2015), a genetic method capable of sparsely labeling individual cells of a neuronal population in multiple colors. 0–1-day old adult flies were heat shocked for 12-13 minutes to induce sparse labeling of LPLC1 or LPLC2.

### Immunohistochemistry

All dissections were performed in ice-cold Schneider’s insect media (S2, Sigma Aldrich, #S01416) to avoid tissue degradation. Brains were then transferred to a 1% paraformaldehyde (20% PFA, Electron Microscopy Sciences, #15713) in S2 solution and fixed overnight at 4°C while rotating. Fixed brains were quickly rinsed 3 times and then washed 4 x 10 min with phosphate buffered saline (pH 7.4) with 0.5% Triton X-100 (PBST) (Sigma-Aldrich, 9002-93-1). Brains were next blocked with 5% Normal Goat Serum (Gibco, 16210064) in PBST (PBST-NGS) in for at least 1.5 hr while rotating at RT. Brains were incubated with primary antibodies in PBST-NGS overnight at 4°C while rotating and washed 3 x 30 min with PBST the next day. Brains were then incubated with secondary antibodies in PBST-NGS overnight at 4°C while rotating, and again washed 3 x 30 min with PBST the next day. Prior to Dylight™ antibody incubation, brains were blocked with 5% Normal Mouse Serum (Invitrogen 31880) in PBST (PBST-NMS) for at least 1.5 hr while rotating at RT. Brains were incubated in DyLight™ antibodies in PBST-NMS overnight at 4°C followed by a minimum of 3 x 30 min PBST washes. After immunostaining, brains were fixed in a 4% PFA in PBS solution for 45 min at RT, followed by 3 x 30 min PBST washes. Brains were then mounted on Poly-L-lysine (Sigma-Aldrich, 25988-63-0) coated glass coverslips (#1.5). Brains were dehydrated in increasing concentrations of ethanol (30%, 50%, 70%, 95%, 100%, 100%) (Decon Labs, 2705HC) for 5 min at each concentration, and then transferred to 100% xylene (Thermofisher scientific, 1330-20-7) for 5 min to clear the brains. Brains were transferred to fresh 100% xylene for another 5 min. Brains were then embedded in DPX (Electron Microscopy Services, 13510) mounting fluid by dropwise application over the affixed brains (∼5-7 drops). The coverslip was then inverted over a microscope slide that had two coverslip spacers (#1) affixed to it with UV glue. The DPX medium was left to cure at RT for at least 24 hr before imaging.

Immunostaining was performed using the following antibodies: mouse anti-nc82 (1:40, DSHB, RRID: AB_2314866), mouse anti-*broad* (1:250, DSHB, RRID: AB_528104), rat anti-DYDDDDK (1:200, Novus Biologicals, RRID: NBP1-06712SS), Alexa Fluor 647 goat anti-mouse (1:400, Thermo Fisher Scientific, RRID: AB_141725), Alexa Fluor 488 goat anti-rat (1:400, Thermo Fisher Scientific, RRID: AB_2534074), 488 anti-HA DyLight™ (1:400, Thermo Fisher Scientific, RRID: AB_2533051), and 550 anti-V5 DyLight™ (1:400, Bio-Rad Antibodies, RRID: MCA1360D550GA).

### Confocal Imaging

All fluorescent images were acquired on an Olympus FV1000 confocal microscope with a 60x, 1.42 NA oil immersion objective (UPlanSApo). For any image quantification, acquisition settings and figure brightness and contrast were kept consistent across samples. Otherwise, parameters for acquisition and figure generation were optimally adjusted.

### Image Analysis

Analyses of morphological data were performed using custom MATLAB scripts. To quantify axonal volume throughout the z-stack of a confocal image, neuronal signal was first thresholded using FIJI’s RenyiEntropy auto thresholding function and then binarized in MATLAB. A region of interest (ROI) was manually drawn and used to restrict axonal quantification to the axons only. The total number of positive pixels was then calculated, and the overall volume was obtained by multiplying the total pixel count by the image voxel size.

To quantify fluorescence within a glomerulus, the middle plane of the glomerulus was first outlined as an ROI according to the Bruchpilot (Brp) stain. Using the ROI, the integrated density signal and area were then taken using the Measure function in Fiji (Schindelin et al., 2012). Corrected total cell fluorescence was then calculated according to the following formula: integrated density – (area * mean fluorescence of background). The Brp stain was also used to visually identify the lobula and lobula plate layers for scoring.

### Connectomics Data Analysis

To determine connectivity partners of LPLC1 and LPLC2 all neurons labeled as LPLC1 (72) and LPLC2 (100) were identified on the right side of the FAFB dataset hosted in the FlyWire codex (version 783; https://codex.flywire.ai/app/search) (Dorkenwald et al., 2022; Matsliah et al., 2024; Schlegel et al., 2023; Zheng et al., 2018). The right side of the brain was selected as the right optic lobe has been extensively characterized, and individual neurons within the fly’s visual system have been identified and proofread (Matsliah et al., 2024). Presynaptic inputs to the entire LPLC1 and LPLC2 populations were retrieved using Python and identified from the FAFB dataset utilizing a confidence metric (cleft_score) of 30 for individual synapses. For each population, a list of synapses, along with coordinate information, presynaptic neuron ID, postsynaptic neuron ID, and cleft_score for each synapse was obtained (Buhmann et al., 2021; Dorkenwald et al., 2022; Heinrich et al., 2018; Zheng et al., 2018). For all analyses of connectivity, only the presynaptic neurons with synapses in the dendritic arbors in the lobula and lobula plate were considered and identified. A total of 30062 unique presynaptic inputs to both the LPLC1 and LPLC2 populations were identified. However, only 29429 of those neurons provided synaptic input to the dendrites in the lobula and lobula plate. Individual neurons that made less than five total synapses to both populations were not identified and were labeled as ‘nan’. Neurons that made five or more synapses but were not identified/labeled in the current version of the codex were labeled as unknown. All other neurons were labeled by their appropriate cell type according to the FlyWire codex (Dorkenwald et al., 2022; Matsliah et al., 2024; Schlegel et al., 2023; Zheng et al., 2018). All neurons labeled as ‘nan’ or unknown were removed from all subsequent analyses. The major synaptic inputs were identified using percentages of total synapses to LPLC1 dendrites by counting the number of synapses a given population made with LPLC1, considering presynaptic populations with a percentage of total inputs >1% as major synaptic inputs for each population. The same methodology was repeated for the LPLC2 population.

For visual comparisons of synapse locations on two individual LPLC1 (FAFB neuron ID: 720575940644654752) and LPLC2 (FAFB neuron ID:720575940611740569), individual mesh reconstructions and synaptic inputs were plotted using R (R version 4.0.5 (*R: A Language and Environment for Statistical Computing*, 2021)). For representative skeleton reconstructions of presynaptic inputs, the following representative example neurons were selected: Tm5f: 720575940608888843, Tm20: 720575940627542018, Tm4: 720575940625995780, T2: 720575940633643033.

### Electrophysiology and Optogenetics

In vivo whole-cell electrophysiology and optogenetic stimulation were performed as described previously (von Reyn et al., 2017). Flies were anesthetized at 4°C with their head and thorax tethered to a polyether ether ketone plate with UV glue (Loctite 3972). The T1 legs were cut at the femur to avoid cleaning of the head. The proboscis was glued in its retracted position to decrease brain movement during the recording. Cuticle and trachea above the LPLC2 population in the right side of the brain were removed and the brain was perfused with standard extracellular saline (NaCl 103 mmol, KCl 3 mmol, TES 5 mmol, trehalose·2H_2_O 8 mmol, glucose 10 mmol, NaHCO_3_ 26 mmol, NaH_2_PO_4_ 1 mmol, CaCl_2_·2H_2_O 1.5 mmol and MgCl_2_·6H_2_O 4 mmol; (Gouwens and Wilson, 2009)). To maintain a pH of 7.3, extracellular saline was adjusted to 270-275 mOsm and bubbled with 95% O_2_/CO_2_. All experiments were performed at room temperature (20-22°C). Localized application of collagenase (0.5% in extracellular saline) with a glass electrode and mechanical pressure was used to break through the brain sheath, providing access to LPLC2 cell bodies. Patch-clamp electrodes (6-9 MΩ) containing intracellular saline (potassium aspartate 140 mmol, KCl 1 mmol, Hepes 10 mmol, EGTA 1 mmol, Na_3_GTP 0.5 mmol, MgATP 4 mmol, Alexafluor-568 5 µmol, 265 mOsm, pH 7.3) were used to target GFP-labeled LPLC2 soma. The membrane voltage was amplified via MultiClamp 700B, digitized (NI-DAQ, National Instruments) at 20 kHz, and low pass-filtered at 6 kHz. Data were obtained using the Wavesurfer (https://wavesurfer.janelia.org/) open-source software running in MATLAB (MathWorks). Recordings were not adjusted for a 13mV liquid junction potential (Gouwens and Wilson, 2009). For recordings to be considered acceptable, an initial seal resistance of >2 GΩ before rupture and a resting membrane potential of −38 mV was required.

For optogenetic experiments, light was delivered (635 nm LED, Scientifica) through a 40x objective. Light activation (100 ms) of T2 cell types expressing CsChrimson was delivered 5 times at 30 second intervals with 3 repetitions, and recordings were taken from LPLC2 cells.

### Electrophysiological Analysis

Analyses of LPLC2 membrane potential peak responses and response latency with respect to the start of the optogenetic light pulse were performed using custom MATLAB scripts. The peak magnitude of the LPLC2 response was measured during the light pulse after baseline (taken one second prior to the light pulse) subtracting the data. Activation latency was measured as the time at which the LPLC2 response exceeded 3 standard deviations of the baseline following optogenetic light stimulation.

### Single-cell Perturb-Seq experiment

Virgin females expressing a nuclear GFP reporter (UAS-His2A-GFP) and either *broad* overexpression constructs or a control (empty) chromosome were crossed to males carrying an LPLC2 split-GAL4 driver (OL0048b), as well as a unique isogenic wild-type X-chromosome from the *Drosophila* Reference Genetic Panel (DRGP) (Huang et al., 2014; Mackay et al., 2012). Each experimental condition was crossed to 3-5 unique DGRP genotypes (see Figure 5 for details). F1 generation females were collected at 0h APF and incubated for either 48h or 72h APF.

All brains were dissected and collected in two Eppendorf tubes kept on ice, one for each timepoint (each timepoint was processed as a separate sample). Brains were incubated in a papain (Worthington #LK003178) and protease (Sigma-Aldrich #5401119001) cocktail at 25°C for 30 min, gently washed twice with PBS, then washed with 0.04% BSA in PBS and dissociated mechanically by pipetting. The resulting cell suspension was filtered through a 20 µm cell-strainer (Corning #352235) and sorted by FACS (BD FACS Aria II) to isolate GFP-positive single cells and measure cell concentrations.

Single-cell suspensions were used to generate scRNA-Seq libraries using the 10X Genomics Chromium Next GEM Single Cell 3’-kit (v3.1) following the manufacturer’s protocol. Each sample (timepoint) was loaded to a single lane of 10X Chromium with a targeted capture rate of 1,500 cells per sample. Two scRNA-Seq libraries were sequenced using one lane of NovaSeq 6000 SP platform (28bp + 91 bp).

### Single-cell RNA-Seq data processing

Raw reads were processed using 10X Cell Ranger (7.1.0) using the reference genome and transcriptome from FlyBase (release 6.29, (Gramates et al., 2022)). The reference genome and transcriptome were appended with the sequence of UAS-His2A-GFP reporter construct to quantify the levels of transgene expression.

Single-cell transcriptomes from individual DGRP-marked biological replicates were demultiplexed based on genotypes of their unique DGRP X-chromosomes using demuxlet (version 2, https://github.com/statgen/popscle; (Kang et al., 2018)). The genotypes of DGRP strains (Huang et al., 2014; Mackay et al., 2012) were preprocessed as described in (Kurmangaliyev et al., 2020). Demultiplexing was based on genetic variants from 18 DGRP strains: 15 strains used in the experiment (not all genotypes yielded high-quality transcriptomes) and 3 strains as negative controls. Variants were filtered using the following criteria: (1) only bi-allelic single-nucleotide polymorphisms (SNP) on X chromosome; (2) SNPs called in all 18 strains (no missing data); (3) non-reference allele only in one of 18 strains. Since we do not have information about the exact genotypes of common maternal chromosomes in F1 heterozygotes, we have quantified allelic coverages of filtered SNPs using samtools mpileup (version 1.10; (Li, 2011)). In heterozygous F1 progeny, the non-reference alleles on a common maternal chromosome are expected to have an allelic frequency of 0.5. Therefore, we only kept SNPs with a minimum coverage of 10 reads and a maximum non-reference allele frequency of 0.25. The resulting set of filtered DGRP SNPs was converted to heterozygous variants using a reference allele as a maternal genotype. In total, 6,765 SNPs were used for demultiplexing. Few cells were incorrectly assigned to the genotypes that were not used in the experiments (negative controls) confirming the accuracy of the demultiplexing step: in the 48h sample, 9 of 1484 cells were assigned to negative controls (244 were “doublets/ambiguous”); in the 72h sample, 18 of 1101 cells were assigned to the negative controls (138 were “doublets/ambiguous”).

### Single-cell RNA-seq data analysis

Single-cell data analysis was performed using Seurat (5.0.1, (Hao et al., 2024)). Each sample (timepoint) was analyzed separately. We kept only high-quality single-cell transcriptomes that were assigned to the expected DGRP genotypes: (UMI/cell: minimum = 10,000, maximum = 50,000; mitochondrial transcripts < 10%). The final datasets included 934 cells for 48h APF and 818 cells for 72h APF. Cells were clustered using the standard Seurat pipeline with default parameters. Total numbers of UMI-per-cell were regressed out at the scaling step (function: ScaleData). Scaled expression values of 2000 highly variable genes were used for principal component analysis, and the first 7 principal components were used for a SNN-based (shared nearest neighbor) clustering (functions: FindNeighbors/FindClusters, resolution = 0.1). The same principal components were used to compute tSNE (t-Distributed Stochastic Neighbor) embeddings. t-SNE plots were used only for the visualization of datasets. Clusters corresponding to LPLC2 neurons were annotated based on the expression of known marker genes (Figure 5).

The transcriptional profiles of LPLC2 neurons from the Perturb-Seq experiment were compared to the transcriptional profiles of LPLC1 and LPLC2 neurons in the single-cell atlas of the developing *Drosophila* visual system (Kurmangaliyev et al., 2020). The comparison was based on differentially expressed genes (DEGs) between LPLC1 and LPLC2 at 48h APF (there were too few LPLC1/LPLC2 cells in the atlas at 72h APF for a meaningful analysis). DEGs were identified using a Wilcoxon rank-sum test (function: FindMarkers, min.pct = 0.5, min.diff.pct = 0.5, pseudocount.use = 0.1). DEGs were filtered using the following criteria: adjusted p-value < 0.01, fold-change > 4, and average normalized expression level > 2 (either in LPLC1 or LPLC2 neurons at 48h APF). The log-scaled average expression profiles of the identified DEGs were compared between LPLC2 neurons in each experimental group in Perturb-Seq and LPLC1/LPLC2 clusters in the atlas using Pearson’s correlation coefficients (Figure 5). Expression patterns of genes of interest were visualized using heatmaps.

The normalized expression values (transcripts-per-10,000, TP10K) were averaged at the levels of cell types, experimental groups, and/or replicates. The average expression values were log1p-transformed and capped at the maximum value of 20 (3 in log-space). The expression of transcription factors in LPLC1 and LPLC2 neurons in Figure 1c is shown for genes with a minimum expression value of 5 in any of the timepoints.

### Isoform analysis of atlas and Perturb-seq data

The differential use of *broad* isoforms was visualized using read coverages at the 3’-UTR region of the gene. Reads corresponding to specific cell types, experimental groups, and time points were extracted from the Cell Ranger output bam-files using Sinto (https://github.com/timoast/sinto), quantified using deepTools2 (Ramirez et al., 2016), and visualized using Integrative genomics viewer (IGV) (Robinson et al., 2011).

### Statistical Analysis

All boxplots were formatted where the dividing line inside the box indicates the median, the grey boxes contain the interquartile range, and the whiskers indicate the data points that fall within 1.5x the interquartile range. Outliers were denoted with a + sign.

Statistical tests were selected based on data distribution (Kolmogorov-Smirnov test, MATLAB) and sample size. All statistical tests are as stated in the figure captions.

## Data and Software Availability

Information about and requests for data can be directed to and will be fulfilled by the Lead Contact, Catherine R. von Reyn. The single-cell transcriptional atlas of the *Drosophila* visual system is available on NCBI GEO (GSE156455) and Zenodo (8111612). The Perturb-Seq dataset will be available on NCBI GEO and Zenodo upon publication.

## Additional Resources

Immunohistochemistry protocols for driver expression visualization and Multi-Color Flip Out: https://www.janelia.org/project-team/flylight/protocols.

## Author Contributions

Conceptualization: N.S., M.D., Y.Z.K., and C.R.v.R.

Data curation: N.S., M.D., B.W.H, A.M-S., J.A., Y.Z.K., and C.R.v.R.

Formal Analysis: N.S., B.W.H, A.M-S., R.G., and Y.Z.K.

Funding acquisition: J.A., M.D. and C.R.v.R.

Investigation: N.S., M.D., B.W.H., A.M-S., R.G., C.R.C., A.R., R.H.H., and P.M.

Methodology: N.S., M.D., Y.Z.K., and C.R.v.R.

Software: N.S., B.W.H., A.M-S., R.G., and Y.Z.K.

Supervision: J.A., Y.Z.K. and C.R.v.R.

Visualization: N.S., B.W.H., A.M-S., Y.Z.K., and C.R.v.R.

Writing – original draft: N.S., Y.Z.K., and C.R.v.R.

Writing – review & editing: N.S., M.D., B.W.H., A.M-S., J.A., Y.Z.K. and C.R.v.R.

## Acknowledgments

We thank all current and past members of the von Reyn and Ausborn laboratories for their discussion of this work. We also thank Larry Zipursky for the use of his laboratory to perform the Perturb-seq experiment. This work was supported by the National Science Foundation Grant No. IOS-1921065 (C.R.v.R.), the National Institutes of Health NINDS R01NS118562 (J.A. & C.R.v.R.), and the HHMI-Helen Hay Whitney Foundation Fellowship (M.D.).

## References

Ache, J. M., Polsky, J., Alghailani, S., Parekh, R., Breads, P., Peek, M. Y., Bock, D. D., von Reyn, C. R., & Card, G. M. (2019). Neural Basis for Looming Size and Velocity Encoding in the Drosophila Giant Fiber Escape Pathway. Curr Biol, 29(6), 1073–1081 e1074. 10.1016/j.cub.2019.01.079

Ahn, S., Yang, H., Son, S., Lee, H. S., Park, D., Yim, H., Choi, H. J., Swoboda, P., & Lee, J. (2022). The C. elegans regulatory factor X (RFX) DAF-19M module: A shift from general ciliogenesis to cell-specific ciliary and behavioral specialization. Cell Rep, 39(2), 110661. 10.1016/j.celrep.2022.110661

Allan, D. W., & Thor, S. (2015). Transcriptional selectors, masters, and combinatorial codes: regulatory principles of neural subtype specification. Wiley Interdiscip Rev Dev Biol, 4(5), 505–528. 10.1002/wdev.191

Arber, S., Han, B., Mendelsohn, M., Smith, M., Jessell, T. M., & Sockanathan, S. (1999). Requirement for the homeobox gene Hb9 in the consolidation of motor neuron identity. Neuron, 23(4), 659–674. 10.1016/s0896-6273(01)80026-x

Bayer, C. A., Holley, B., & Fristrom, J. W. (1996). A switch in broad-complex zinc-finger isoform expression is regulated posttranscriptionally during the metamorphosis of Drosophila imaginal discs. Dev Biol, 177(1), 1–14. 10.1006/dbio.1996.0140

Bayer, C. A., von Kalm, L., & Fristrom, J. W. (1997). Relationships between protein isoforms and genetic functions demonstrate functional redundancy at the Broad-Complex during Drosophila metamorphosis. Dev Biol, 187(2), 267–282. 10.1006/dbio.1997.8620

Belyaeva, E. S., Aizenzon, M. G., Semeshin, V. F., Kiss, II, Koczka, K., Baritcheva, E. M., Gorelova, T. D., & Zhimulev, I. F. (1980). Cytogenetic analysis of the 2B3-4--2B11 region of the X-chromosome of Drosophila melanogaster. I. Cytology of the region and mutant complementation groups. Chromosoma, 81(2), 281–306. 10.1007/BF00285954

Brennan, C. A., Li, T. R., Bender, M., Hsiung, F., & Moses, K. (2001). Broad-complex, but not ecdysone receptor, is required for progression of the morphogenetic furrow in the Drosophila eye. Development, 128(1), 1–11. 10.1242/dev.128.1.1

Buhmann, J., Sheridan, A., Malin-Mayor, C., Schlegel, P., Gerhard, S., Kazimiers, T., Krause, R., Nguyen, T. M., Heinrich, L., Lee, W. A., Wilson, R., Saalfeld, S., Jefferis, G., Bock, D. D., Turaga, S. C., Cook, M., & Funke, J. (2021). Automatic detection of synaptic partners in a whole-brain Drosophila electron microscopy data set. Nat Methods, 18(7), 771–774. 10.1038/s41592-021-01183-7

Campbell, R. F., & Walthall, W. W. (2016). Meis/UNC-62 isoform dependent regulation of CoupTF-II/UNC-55 and GABAergic motor neuron subtype differentiation. Dev Biol, 419(2), 250–261. 10.1016/j.ydbio.2016.09.009

Chao, H. T., Davids, M., Burke, E., Pappas, J. G., Rosenfeld, J. A., McCarty, A. J., Davis, T., Wolfe, L., Toro, C., Tifft, C., Xia, F., Stong, N., Johnson, T. K., Warr, C. G., Undiagnosed Diseases, N., Yamamoto, S., Adams, D. R., Markello, T. C., Gahl, W. A., Bellen, H. J., Wangler, M. F., & Malicdan, M. C. V. (2017). A Syndromic Neurodevelopmental Disorder Caused by De Novo Variants in EBF3. Am J Hum Genet, 100(1), 128–137. 10.1016/j.ajhg.2016.11.018

Chen, Y. C., & Konstantinides, N. (2022). Integration of Spatial and Temporal Patterning in the Invertebrate and Vertebrate Nervous System. Front Neurosci, 16, 854422. 10.3389/fnins.2022.854422

Chu, S. Y., Lai, Y. W., Hsu, T. C., Lu, T. M., & Yu, H. H. (2024). Isoforms of Terminal Selector Mamo Control Axon Segregation During Adult Drosophila Memory Center Construction Via Semaphorin-1a. 10.2139/ssrn.4749745

Crossgrove, K., Bayer, C. A., Fristrom, J. W., & Guild, G. M. (1996). The Drosophila Broad-Complex early gene directly regulates late gene transcription during the ecdysone-induced puffing cascade. Dev Biol, 180(2), 745–758. 10.1006/dbio.1996.0343

Davis, F. P., Nern, A., Picard, S., Reiser, M. B., Rubin, G. M., Eddy, S. R., & Henry, G. L. (2020). A genetic, genomic, and computational resource for exploring neural circuit function. Elife, 9. 10.7554/eLife.50901

Deneris, E. S., & Hobert, O. (2014). Maintenance of postmitotic neuronal cell identity. Nat Neurosci, 17(7), 899–907. 10.1038/nn.3731

DiBello, P. R., Withers, D. A., Bayer, C. A., Fristrom, J. W., & Guild, G. M. (1991). The Drosophila Broad-Complex encodes a family of related proteins containing zinc fingers. Genetics, 129(2), 385–397. 10.1093/genetics/129.2.385

Dionne, H., Hibbard, K. L., Cavallaro, A., Kao, J. C., & Rubin, G. M. (2018). Genetic Reagents for Making Split-GAL4 Lines in Drosophila. Genetics, 209(1), 31–35. 10.1534/genetics.118.300682

Dorkenwald, S., McKellar, C. E., Macrina, T., Kemnitz, N., Lee, K., Lu, R., Wu, J., Popovych, S., Mitchell, E., Nehoran, B., Jia, Z., Bae, J. A., Mu, S., Ih, D., Castro, M., Ogedengbe, O., Halageri, A., Kuehner, K., Sterling, A. R., Ashwood, Z., Zung, J., Brittain, D., Collman, F., Schneider-Mizell, C., Jordan, C., Silversmith, W., Baker, C., Deutsch, D., Encarnacion-Rivera, L., Kumar, S., Burke, A., Bland, D., Gager, J., Hebditch, J., Koolman, S., Moore, M., Morejohn, S., Silverman, B., Willie, K., Willie, R., Yu, S. C., Murthy, M., & Seung, H. S. (2022). FlyWire: online community for whole-brain connectomics. Nat Methods, 19(1), 119–128. 10.1038/s41592-021-01330-0

El-Danaf, R. N., Rajesh, R., & Desplan, C. (2023). Temporal regulation of neural diversity in Drosophila and vertebrates. Semin Cell Dev Biol, 142, 13–22. 10.1016/j.semcdb.2022.05.011

Emery, I. F., Bedian, V., & Guild, G. M. (1994). Differential expression of Broad-Complex transcription factors may forecast tissue-specific developmental fates during Drosophila metamorphosis. Development, 120(11), 3275–3287. 10.1242/dev.120.11.3275

Fischbach, K. F., & Dittrich, A. P. (1989). The optic lobe of Drosophila melanogaster. I. A Golgi analysis of wild-type structure. Cell and Tissue Research, 258, 441–475. 10.1007/BF00218858

Flames, N., & Hobert, O. (2009). Gene regulatory logic of dopamine neuron differentiation. Nature, 458(7240), 885–889. 10.1038/nature07929

Gramates, L. S., Agapite, J., Attrill, H., Calvi, B. R., Crosby, M. A., Dos Santos, G., Goodman, J. L., Goutte-Gattat, D., Jenkins, V. K., Kaufman, T., Larkin, A., Matthews, B. B., Millburn, G., Strelets, V. B., & the FlyBase, C. (2022). FlyBase: a guided tour of highlighted features. Genetics, 220(4). 10.1093/genetics/iyac035

Hao, Y., Stuart, T., Kowalski, M. H., Choudhary, S., Hoffman, P., Hartman, A., Srivastava, A., Molla, G., Madad, S., Fernandez-Granda, C., & Satija, R. (2024). Dictionary learning for integrative, multimodal and scalable single-cell analysis. Nat Biotechnol, 42(2), 293–304. 10.1038/s41587-023-01767-y

Hassan, B. A., & Hiesinger, P. R. (2015). Beyond Molecular Codes: Simple Rules to Wire Complex Brains. Cell, 163(2), 285–291. 10.1016/j.cell.2015.09.031

Heinrich, L., Funke, J., Pape, C., Nunez-Iglesias, J., & Saalfeld, S. (2018). Synaptic Cleft Segmentation in Non-isotropic Volume Electron Microscopy of the Complete Drosophila Brain. MICCAI 2018,

Hobert, O. (2008). Regulatory logic of neuronal diversity: Terminal selector genes and selector motifs. Proceedings of the National Academy of Sciences, 105(51), 20067–20071. 10.1073/pnas.0806070105

Hobert, O. (2016). Terminal Selectors of Neuronal Identity. Curr Top Dev Biol, 116, 455–475. 10.1016/bs.ctdb.2015.12.007

Hobert, O. (2021). Homeobox genes and the specification of neuronal identity. Nat Rev Neurosci, 22(10), 627–636. 10.1038/s41583-021-00497-x

Hobert, O., & Kratsios, P. (2019). Neuronal identity control by terminal selectors in worms, flies, and chordates. Curr Opin Neurobiol, 56, 97–105. 10.1016/j.conb.2018.12.006

Holguera, I., & Desplan, C. (2018). Neuronal specification in space and time. Science, 362(6411), 176–180. 10.1126/science.aas9435

Holmberg, J., & Perlmann, T. (2012). Maintaining differentiated cellular identity. Nat Rev Genet, 13(6), 429–439. 10.1038/nrg3209

Huang, W., Massouras, A., Inoue, Y., Peiffer, J., Ramia, M., Tarone, A. M., Turlapati, L., Zichner, T., Zhu, D., Lyman, R. F., Magwire, M. M., Blankenburg, K., Carbone, M. A., Chang, K., Ellis, L. L., Fernandez, S., Han, Y., Highnam, G., Hjelmen, C. E., Jack, J. R., Javaid, M., Jayaseelan, J., Kalra, D., Lee, S., Lewis, L., Munidasa, M., Ongeri, F., Patel, S., Perales, L., Perez, A., Pu, L., Rollmann, S. M., Ruth, R., Saada, N., Warner, C., Williams, A., Wu, Y. Q., Yamamoto, A., Zhang, Y., Zhu, Y., Anholt, R. R., Korbel, J. O., Mittelman, D., Muzny, D. M., Gibbs, R. A., Barbadilla, A., Johnston, J. S., Stone, E. A., Richards, S., Deplancke, B., & Mackay, T. F. (2014). Natural variation in genome architecture among 205 Drosophila melanogaster Genetic Reference Panel lines. Genome Res, 24(7), 1193–1208. 10.1101/gr.171546.113

Huet, F., Ruiz, C., & Richards, G. (1993). Puffs and PCR: the in vivo dynamics of early gene expression during ecdysone responses in Drosophila. Development, 118(2), 613–627. 10.1242/dev.118.2.613

Jain, S., Lin, Y., Kurmangaliyev, Y. Z., Valdes-Aleman, J., LoCascio, S. A., Mirshahidi, P., Parrington, B., & Zipursky, S. L. (2022). A global timing mechanism regulates cell-type-specific wiring programmes. Nature, 603(7899), 112–118. 10.1038/s41586-022-04418-5

Jenett, A., Rubin, G. M., Ngo, T. T., Shepherd, D., Murphy, C., Dionne, H., Pfeiffer, B. D., Cavallaro, A., Hall, D., Jeter, J., Iyer, N., Fetter, D., Hausenfluck, J. H., Peng, H., Trautman, E. T., Svirskas, R. R., Myers, E. W., Iwinski, Z. R., Aso, Y., DePasquale, G. M., Enos, A., Hulamm, P., Lam, S. C., Li, H. H., Laverty, T. R., Long, F., Qu, L., Murphy, S. D., Rokicki, K., Safford, T., Shaw, K., Simpson, J. H., Sowell, A., Tae, S., Yu, Y., & Zugates, C. T. (2012). A GAL4-driver line resource for Drosophila neurobiology. Cell Rep, 2(4), 991–1001. 10.1016/j.celrep.2012.09.011

Kang, H. M., Subramaniam, M., Targ, S., Nguyen, M., Maliskova, L., McCarthy, E., Wan, E., Wong, S., Byrnes, L., Lanata, C. M., Gate, R. E., Mostafavi, S., Marson, A., Zaitlen, N., Criswell, L. A., & Ye, C. J. (2018). Multiplexed droplet single-cell RNA-sequencing using natural genetic variation. Nat Biotechnol, 36(1), 89–94. 10.1038/nbt.4042

Karim, F. D., Guild, G. M., & Thummel, C. S. (1993). The Drosophila Broad-Complex plays a key role in controlling ecdysone-regulated gene expression at the onset of metamorphosis. Development, 118(3), 977–988. 10.1242/dev.118.3.977

Kiss, I., Beaton, A. H., Tardiff, J., Fristrom, D., & Fristrom, J. W. (1988). Interactions and developmental effects of mutations in the Broad-Complex of Drosophila melanogaster. Genetics, 118(2), 247–259. 10.1093/genetics/118.2.247

Kiss, I., Bencze, G., Fodor, G., Szabad, J., & Fristrom, J. W. (1976). Prepupal larval mosaics in Drosophila melanogaster. Nature, 262(5564), 136–138. 10.1038/262136a0

Klapoetke, N. C., Nern, A., Peek, M. Y., Rogers, E. M., Breads, P., Rubin, G. M., Reiser, M. B., & Card, G. M. (2017). Ultra-selective looming detection from radial motion opponency. Nature, 551(7679), 237–241. 10.1038/nature24626

Klapoetke, N. C., Nern, A., Rogers, E. M., Rubin, G. M., Reiser, M. B., & Card, G. M. (2022). A functionally ordered visual feature map in the Drosophila brain. Neuron. 10.1016/j.neuron.2022.02.013

Konstantinides, N., Kapuralin, K., Fadil, C., Barboza, L., Satija, R., & Desplan, C. (2018). Phenotypic Convergence: Distinct Transcription Factors Regulate Common Terminal Features. Cell, 174(3), 622–635 e613. 10.1016/j.cell.2018.05.021

Kumar, J. P. (2001). Signalling pathways in Drosophila and vertebrate retinal development. Nat Rev Genet, 2(11), 846–857. 10.1038/35098564

Kurmangaliyev, Y. Z., Yoo, J., Valdes-Aleman, J., Sanfilippo, P., & Zipursky, S. L. (2020). Transcriptional Programs of Circuit Assembly in the Drosophila Visual System. Neuron, 108(6), 1045–1057 e1046. 10.1016/j.neuron.2020.10.006

Lai, S. L., & Lee, T. (2006). Genetic mosaic with dual binary transcriptional systems in Drosophila. Nat Neurosci, 9(5), 703–709. 10.1038/nn1681

Li, H. (2011). A statistical framework for SNP calling, mutation discovery, association mapping and population genetical parameter estimation from sequencing data. Bioinformatics, 27(21), 2987–2993. 10.1093/bioinformatics/btr509

Li, X., Erclik, T., Bertet, C., Chen, Z., Voutev, R., Venkatesh, S., Morante, J., Celik, A., & Desplan, C. (2013). Temporal patterning of Drosophila medulla neuroblasts controls neural fates. Nature, 498(7455), 456–462. 10.1038/nature12319

Mackay, T. F., Richards, S., Stone, E. A., Barbadilla, A., Ayroles, J. F., Zhu, D., Casillas, S., Han, Y., Magwire, M. M., Cridland, J. M., Richardson, M. F., Anholt, R. R., Barron, M., Bess, C., Blankenburg, K. P., Carbone, M. A., Castellano, D., Chaboub, L., Duncan, L., Harris, Z., Javaid, M., Jayaseelan, J. C., Jhangiani, S. N., Jordan, K. W., Lara, F., Lawrence, F., Lee, S. L., Librado, P., Linheiro, R. S., Lyman, R. F., Mackey, A. J., Munidasa, M., Muzny, D. M., Nazareth, L., Newsham, I., Perales, L., Pu, L. L., Qu, C., Ramia, M., Reid, J. G., Rollmann, S. M., Rozas, J., Saada, N., Turlapati, L., Worley, K. C., Wu, Y. Q., Yamamoto, A., Zhu, Y., Bergman, C. M., Thornton, K. R., Mittelman, D., & Gibbs, R. A. (2012). The Drosophila melanogaster Genetic Reference Panel. Nature, 482(7384), 173–178. 10.1038/nature10811

Makos, M. A., Kuklinski, N. J., Berglund, E. C., Heien, M. L., & Ewing, A. G. (2009). Chemical measurements in Drosophila. Trends Analyt Chem, 28(11), 1223–1234. 10.1016/j.trac.2009.08.005

Matsliah, Szi-chieh Yu, Krzysztof Kruk, Doug Bland, Austin Burke, Jay Gager, James Hebditch, Ben Silverman, Kyle Willie, Ryan Willie, Marissa Sorek, Amy R. Sterling, Emil Kind, Dustin Garner, Gizem Sancer, Mathias F. Wernet, Sung Soo Kim, Mala Murthy, H. Sebastian Seung, & Consortium, t. F. (2024). Neuronal “parts list” and wiring diagram for a visual system. 10.1101/2023.10.12.562119

McFarland, B. W., Smolin, N., Jang, H., Hina, B. W., Parisi, M. J., Davis, K. C., Mosca, T. J., Godenschewege, T. A., Nern, A., Kurmangaliyev, Y. Z., & von Reyn, C. R. (2024). Axon arrival times and physical occupancy establish visual projection neuron integration on developing dendrites in the Drosophila optic glomeruli. Elife, 13:RP96223. 10.7554/eLife.96223.1

Miller, J., McLachlan, A. D., & Klug, A. (1985). Repetitive zinc-binding domains in the protein transcription factor IIIA from Xenopus oocytes. EMBO J, 4(6), 1609–1614. 10.1002/j.1460-2075.1985.tb03825.x

Mugat, B., Brodu, V., Kejzlarova-Lepesant, J., Antoniewski, C., Bayer, C. A., Fristrom, J. W., & Lepesant, J. A. (2000). Dynamic expression of broad-complex isoforms mediates temporal control of an ecdysteroid target gene at the onset of Drosophila metamorphosis. Dev Biol, 227(1), 104–117. 10.1006/dbio.2000.9879

Nern, A., Loesche, F., Takemura, S. Y., Burnett, L. E., Dreher, M., Gruntman, E., Hoeller, J., Huang, G. B., Januszewski, M., Klapoetke, N. C., Koskela, S., Longden, K. D., Lu, Z., Preibisch, S., Qiu, W., Rogers, E. M., Seenivasan, P., Zhao, A., Bogovic, J., Canino, B. S., Clements, J., Cook, M., Finley-May, S., Flynn, M. A., Fragniere, A. M., Hameed, I., Hayworth, K. J., Hopkins, G. P., Hubbard, P. M., Katz, W. T., Kovalyak, J., Lauchie, S. A., Leonard, M., Lohff, A., Maldonado, C. A., Mooney, C., Okeoma, N., Olbris, D. J., Ordish, C., Paterson, T., Phillips, E. M., Pietzsch, T., Rivas Salinas, J., Rivlin, P. K., Schlegel, P., Scott, A. L., Scuderi, L. A., Takemura, S., Talebi, I., Thomson, A., Trautman, E. T., Umayam, L., Walsh, C., Walsh, J. J., Xu, C. S., Yakal, E. A., Yang, T., Zhao, T., Funke, J., George, R., Hess, H. F., Jefferis, G., Knecht, C., Korff, W., Plaza, S. M., Romani, S., Saalfeld, S., Scheffer, L. K., Berg, S., Rubin, G. M., & Reiser, M. B. (2024). Connectome-driven neural inventory of a complete visual system. bioRxiv. 10.1101/2024.04.16.589741

Nern, A., Pfeiffer, B. D., & Rubin, G. M. (2015). Optimized tools for multicolor stochastic labeling reveal diverse stereotyped cell arrangements in the fly visual system. Proc Natl Acad Sci U S A, 112(22), E2967–2976. 10.1073/pnas.1506763112

Neville, M. C., Nojima, T., Ashley, E., Parker, D. J., Walker, J., Southall, T., Van de Sande, B., Marques, A. C., Fischer, B., Brand, A. H., Russell, S., Ritchie, M. G., Aerts, S., & Goodwin, S. F. (2014). Male-specific fruitless isoforms target neurodevelopmental genes to specify a sexually dimorphic nervous system. Curr Biol, 24(3), 229–241. 10.1016/j.cub.2013.11.035

O’Meara, M. M., Zhang, F., & Hobert, O. (2010). Maintenance of neuronal laterality in Caenorhabditis elegans through MYST histone acetyltransferase complex components LSY-12, LSY-13 and LIN-49. Genetics, 186(4), 1497–1502. 10.1534/genetics.110.123661

Ozel, M. N., Gibbs, C. S., Holguera, I., Soliman, M., Bonneau, R., & Desplan, C. (2022). Coordinated control of neuronal differentiation and wiring by sustained transcription factors. Science, 378(6626), eadd1884. 10.1126/science.add1884

Ozel, M. N., Simon, F., Jafari, S., Holguera, I., Chen, Y. C., Benhra, N., El-Danaf, R. N., Kapuralin, K., Malin, J. A., Konstantinides, N., & Desplan, C. (2021). Neuronal diversity and convergence in a visual system developmental atlas. Nature, 589(7840), 88–95. 10.1038/s41586-020-2879-3

Pereira, L., Aeschimann, F., Wang, C., Lawson, H., Serrano-Saiz, E., Portman, D. S., Grosshans, H., & Hobert, O. (2019). Timing mechanism of sexually dimorphic nervous system differentiation. Elife, 8. 10.7554/eLife.42078

Perkins, L. A., Holderbaum, L., Tao, R., Hu, Y., Sopko, R., McCall, K., Yang-Zhou, D., Flockhart, I., Binari, R., Shim, H. S., Miller, A., Housden, A., Foos, M., Randkelv, S., Kelley, C., Namgyal, P., Villalta, C., Liu, L. P., Jiang, X., Huan-Huan, Q., Wang, X., Fujiyama, A., Toyoda, A., Ayers, K., Blum, A., Czech, B., Neumuller, R., Yan, D., Cavallaro, A., Hibbard, K., Hall, D., Cooley, L., Hannon, G. J., Lehmann, R., Parks, A., Mohr, S. E., Ueda, R., Kondo, S., Ni, J. Q., & Perrimon, N. (2015). The Transgenic RNAi Project at Harvard Medical School: Resources and Validation. Genetics, 201(3), 843–852. 10.1534/genetics.115.180208

Pfeiffer, B. D., Ngo, T. T., Hibbard, K. L., Murphy, C., Jenett, A., Truman, J. W., & Rubin, G. M. (2010). Refinement of tools for targeted gene expression in Drosophila. Genetics, 186(2), 735–755. 10.1534/genetics.110.119917

R: A Language and Environment for Statistical Computing. In. (2021). R Foundation for Statistical Computing. https://www.R-project.org/

Ramirez, F., Ryan, D. P., Gruning, B., Bhardwaj, V., Kilpert, F., Richter, A. S., Heyne, S., Dundar, F., & Manke, T. (2016). deepTools2: a next generation web server for deep-sequencing data analysis. Nucleic Acids Res, 44(W1), W160–165. 10.1093/nar/gkw257

Remesal, L., Roger-Baynat, I., Chirivella, L., Maicas, M., Brocal-Ruiz, R., Perez-Villalba, A., Cucarella, C., Casado, M., & Flames, N. (2020). PBX1 acts as terminal selector for olfactory bulb dopaminergic neurons. Development, 147(8). 10.1242/dev.186841

Restifo, L. L., & White, K. (1991). Mutations in a steroid hormone-regulated gene disrupt the metamorphosis of the central nervous system in Drosophila. Dev Biol, 148(1), 174–194. 10.1016/0012-1606(91)90328-z

Robinson, J. T., Thorvaldsdottir, H., Winckler, W., Guttman, M., Lander, E. S., Getz, G., & Mesirov, J. P. (2011). Integrative genomics viewer. Nat Biotechnol, 29(1), 24–26. 10.1038/nbt.1754

Sahay, A., & Hen, R. (2007). Adult hippocampal neurogenesis in depression. Nat Neurosci, 10(9), 1110–1115. 10.1038/nn1969

Scheffer, L. K., Xu, C. S., Januszewski, M., Lu, Z., Takemura, S. Y., Hayworth, K. J., Huang, G. B., Shinomiya, K., Maitlin-Shepard, J., Berg, S., Clements, J., Hubbard, P. M., Katz, W. T., Umayam, L., Zhao, T., Ackerman, D., Blakely, T., Bogovic, J., Dolafi, T., Kainmueller, D., Kawase, T., Khairy, K. A., Leavitt, L., Li, P. H., Lindsey, L., Neubarth, N., Olbris, D. J., Otsuna, H., Trautman, E. T., Ito, M., Bates, A. S., Goldammer, J., Wolff, T., Svirskas, R., Schlegel, P., Neace, E., Knecht, C. J., Alvarado, C. X., Bailey, D. A., Ballinger, S., Borycz, J. A., Canino, B. S., Cheatham, N., Cook, M., Dreher, M., Duclos, O., Eubanks, B., Fairbanks, K., Finley, S., Forknall, N., Francis, A., Hopkins, G. P., Joyce, E. M., Kim, S., Kirk, N. A., Kovalyak, J., Lauchie, S. A., Lohff, A., Maldonado, C., Manley, E. A., McLin, S., Mooney, C., Ndama, M., Ogundeyi, O., Okeoma, N., Ordish, C., Padilla, N., Patrick, C. M., Paterson, T., Phillips, E. E., Phillips, E. M., Rampally, N., Ribeiro, C., Robertson, M. K., Rymer, J. T., Ryan, S. M., Sammons, M., Scott, A. K., Scott, A. L., Shinomiya, A., Smith, C., Smith, K., Smith, N. L., Sobeski, M. A., Suleiman, A., Swift, J., Takemura, S., Talebi, I., Tarnogorska, D., Tenshaw, E., Tokhi, T., Walsh, J. J., Yang, T., Horne, J. A., Li, F., Parekh, R., Rivlin, P. K., Jayaraman, V., Costa, M., Jefferis, G. S., Ito, K., Saalfeld, S., George, R., Meinertzhagen, I. A., Rubin, G. M., Hess, H. F., Jain, V., & Plaza, S. M. (2020). A connectome and analysis of the adult Drosophila central brain. Elife, 9. 10.7554/eLife.57443

Schindelin, J., Arganda-Carreras, I., Frise, E., Kaynig, V., Longair, M., Pietzsch, T., Preibisch, S., Rueden, C., Saalfeld, S., Schmid, B., Tinevez, J. Y., White, D. J., Hartenstein, V., Eliceiri, K., Tomancak, P., & Cardona, A. (2012). Fiji: an open-source platform for biological-image analysis. Nat Methods, 9(7), 676–682. 10.1038/nmeth.2019

Schizophrenia Working Group of the Psychiatric Genomics, C. (2014). Biological insights from 108 schizophrenia-associated genetic loci. Nature, 511(7510), 421–427. 10.1038/nature13595

Schlegel, P., Yin, Y., Bates, A. S., Dorkenwald, S., Eichler, K., Brooks, P., Han, D. S., Gkantia, M., Dos Santos, M., Munnelly, E. J., Badalamente, G., Capdevila, L. S., Sane, V. A., Pleijzier, M. W., Tamimi, I. F. M., Dunne, C. R., Salgarella, I., Javier, A., Fang, S., Perlman, E., Kazimiers, T., Jagannathan, S. R., Matsliah, A., Sterling, A. R., Yu, S. C., McKellar, C. E., FlyWire, C., Costa, M., Seung, H. S., Murthy, M., Hartenstein, V., Bock, D. D., & Jefferis, G. (2023). Whole-brain annotation and multi-connectome cell typing quantifies circuit stereotypy in Drosophila. bioRxiv. 10.1101/2023.06.27.546055

Serrano-Saiz, E., Poole, R. J., Felton, T., Zhang, F., De La Cruz, E. D., & Hobert, O. (2013). Modular control of glutamatergic neuronal identity in C. elegans by distinct homeodomain proteins. Cell, 155(3), 659–673. 10.1016/j.cell.2013.09.052

Sleven, H., Welsh, S. J., Yu, J., Churchill, M. E. A., Wright, C. F., Henderson, A., Horvath, R., Rankin, J., Vogt, J., Magee, A., McConnell, V., Green, A., King, M. D., Cox, H., Armstrong, L., Lehman, A., Nelson, T. N., Deciphering Developmental Disorders, s., study, C., Williams, J., Clouston, P., Hagman, J., & Nemeth, A. H. (2017). De Novo Mutations in EBF3 Cause a Neurodevelopmental Syndrome. Am J Hum Genet, 100(1), 138–150. 10.1016/j.ajhg.2016.11.020

Takemura, S. Y., Bharioke, A., Lu, Z., Nern, A., Vitaladevuni, S., Rivlin, P. K., Katz, W. T., Olbris, D. J., Plaza, S. M., Winston, P., Zhao, T., Horne, J. A., Fetter, R. D., Takemura, S., Blazek, K., Chang, L. A., Ogundeyi, O., Saunders, M. A., Shapiro, V., Sigmund, C., Rubin, G. M., Scheffer, L. K., Meinertzhagen, I. A., & Chklovskii, D. B. (2013). A visual motion detection circuit suggested by Drosophila connectomics. Nature, 500(7461), 175–181. 10.1038/nature12450

Takemura, S. Y., Xu, C. S., Lu, Z., Rivlin, P. K., Parag, T., Olbris, D. J., Plaza, S., Zhao, T., Katz, W. T., Umayam, L., Weaver, C., Hess, H. F., Horne, J. A., Nunez-Iglesias, J., Aniceto, R., Chang, L. A., Lauchie, S., Nasca, A., Ogundeyi, O., Sigmund, C., Takemura, S., Tran, J., Langille, C., Le Lacheur, K., McLin, S., Shinomiya, A., Chklovskii, D. B., Meinertzhagen, I. A., & Scheffer, L. K. (2015). Synaptic circuits and their variations within different columns in the visual system of Drosophila. Proc Natl Acad Sci U S A, 112(44), 13711–13716. 10.1073/pnas.1509820112

Tanaka, R., & Clark, D. A. (2022). Neural mechanisms to exploit positional geometry for collision avoidance. Curr Biol, 32(11), 2357–2374 e2356. 10.1016/j.cub.2022.04.023

Tsuchida, T., Ensini, M., Morton, S. B., Baldassare, M., Edlund, T., Jessell, T. M., & Pfaff, S. L. (1994). Topographic organization of embryonic motor neurons defined by expression of LIM homeobox genes. Cell, 79(6), 957–970. 10.1016/0092-8674(94)90027-2

von Reyn, C. R., Breads, P., Peek, M. Y., Zheng, G. Z., Williamson, W. R., Yee, A. L., Leonardo, A., & Card, G. M. (2014). A spike-timing mechanism for action selection. Nat Neurosci, 17(7), 962–970. 10.1038/nn.3741

von Reyn, C. R., Nern, A., Williamson, W. R., Breads, P., Wu, M., Namiki, S., & Card, G. M. (2017). Feature Integration Drives Probabilistic Behavior in the Drosophila Escape Response. Neuron, 94(6), 1190–1204 e1196. 10.1016/j.neuron.2017.05.036

Wolfram, V., Southall, T. D., Gunay, C., Prinz, A. A., Brand, A. H., & Baines, R. A. (2014). The transcription factors islet and Lim3 combinatorially regulate ion channel gene expression. J Neurosci, 34(7), 2538–2543. 10.1523/JNEUROSCI.4511-13.2014

Wu, M., Nern, A., Williamson, W. R., Morimoto, M. M., Reiser, M. B., Card, G. M., & Rubin, G. M. (2016). Visual projection neurons in the Drosophila lobula link feature detection to distinct behavioral programs. Elife, 5. 10.7554/eLife.21022

Wyler, S. C., Spencer, W. C., Green, N. H., Rood, B. D., Crawford, L., Craige, C., Gresch, P., McMahon, D. G., Beck, S. G., & Deneris, E. (2016). Pet-1 Switches Transcriptional Targets Postnatally to Regulate Maturation of Serotonin Neuron Excitability. J Neurosci, 36(5), 1758–1774. 10.1523/JNEUROSCI.3798-15.2016

Yao, C., Sasaki, H. M., Ueda, T., Tomari, Y., & Tadakuma, H. (2015). Single-Molecule Analysis of the Target Cleavage Reaction by the Drosophila RNAi Enzyme Complex. Mol Cell, 59(1), 125–132. 10.1016/j.molcel.2015.05.015

Zheng, Z., Lauritzen, J. S., Perlman, E., Robinson, C. G., Nichols, M., Milkie, D., Torrens, O., Price, J., Fisher, C. B., Sharifi, N., Calle-Schuler, S. A., Kmecova, L., Ali, I. J., Karsh, B., Trautman, E. T., Bogovic, J. A., Hanslovsky, P., Jefferis, G., Kazhdan, M., Khairy, K., Saalfeld, S., Fetter, R. D., & Bock, D. D. (2018). A Complete Electron Microscopy Volume of the Brain of Adult Drosophila melanogaster. Cell, 174(3), 730–743 e722. 10.1016/j.cell.2018.06.019

Zhou, B., Williams, D. W., Altman, J., Riddiford, L. M., & Truman, J. W. (2009). Temporal patterns of broad isoform expression during the development of neuronal lineages in Drosophila. Neural Dev, 4, 39. 10.1186/1749-8104-4-39

