## Supplemental Figures for "Neuronal identity control at the resolution of a single transcription factor isoform"

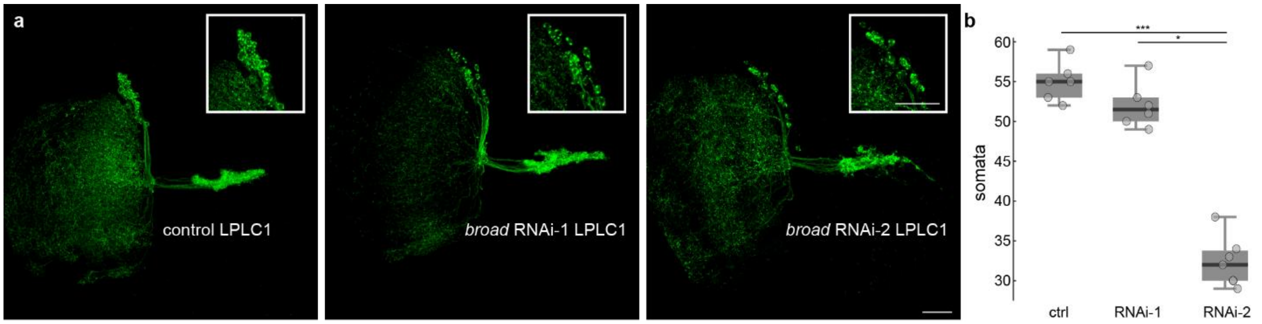


**Supplementary Figure 1: Decreased number of somata labeled in *broad* RNAi-2 LPLC1 cells.**

**(a)** Max projection of control LPLC1, *broad* RNAi-1 LPLC1, and *broad* RNAi-2 LPLC1 populations. Scale bar = 20 µm. Inset: magnified cell body populations. Inset scale bar = 20 µm.

**(b)** Quantification of somata within each group. N ≥ 4 animals for each condition. Kruskal Wallis (p = 1 9.48e-04) Dunn-Sidak post hoc * = p<0.05, ** = p<0.01, *** = p<0.001.

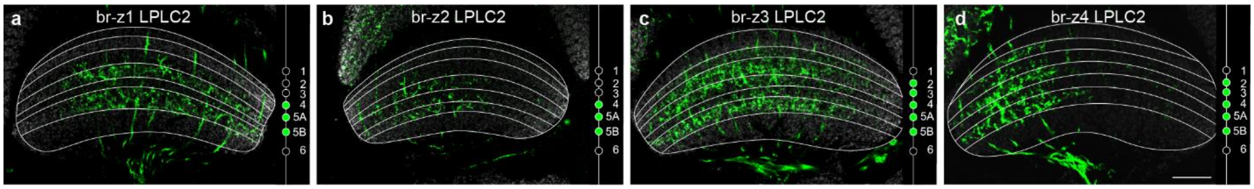


**Supplementary Figure 2: Only *broad-z3* and *-z4* alter dendrite innervation patterns of LPLC2 cells.**

**(a-d)** Single plane LPLC2 dendritic innervations of the lobula for **(a)** *broad-z1* cells, **(b)** *broad-z2* cells, **(c)** *broad-z3* cells, and **(d)** *broad-z4* cells. N ≥ 4 animals for each condition. Scale bar = 20 µm.

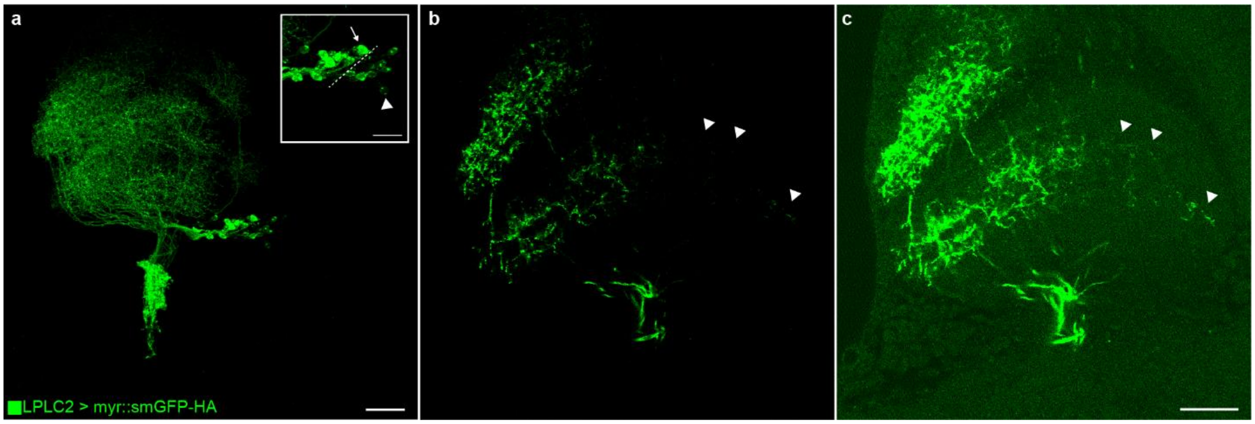


**Supplementary Figure 3: *broad-z4* overexpression results in weakly labeled LPLC2 cells.**

**(a)** Max projection of *broad-z4* LPLC2 cells. Inset: Two distinct populations of cells appear, one brightly labeled (arrow) and one dimly labeled (arrowhead). Scale bar = 10 µm.

**(b-c)** Dorsal loss of dendrites occur upon **(b)** *broad-z4* overexpression. Brightness and contrast adjusted images **(c)** reveal weakly labeled dendrites (arrowheads). Scale bar = 20 µm.

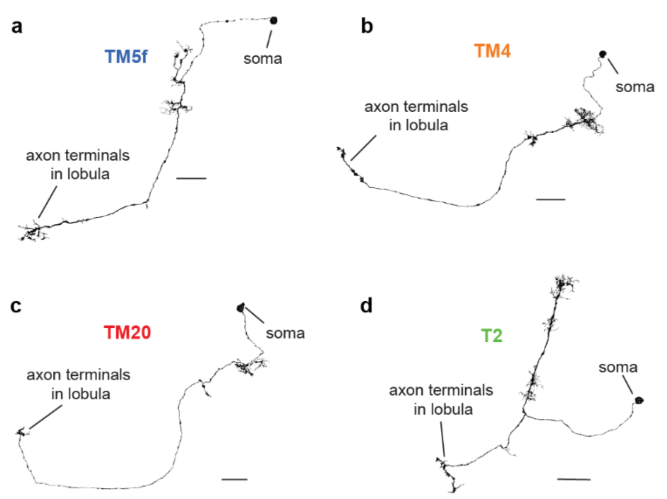


**Supplementary Figure 5: Mesh reconstructions for representative examples of the major LPLC1 and LPLC2 inputs.**

**(a-d)** Mesh reconstructions of **(a)** TM5f, **(b)** TM4, **(c)** TM20, and **(d)** T2. **(a-c)** are the top shared inputs to LPLC1 and LPLC2 and **(d)** is the top differential input. Scalebars = 15 μm.

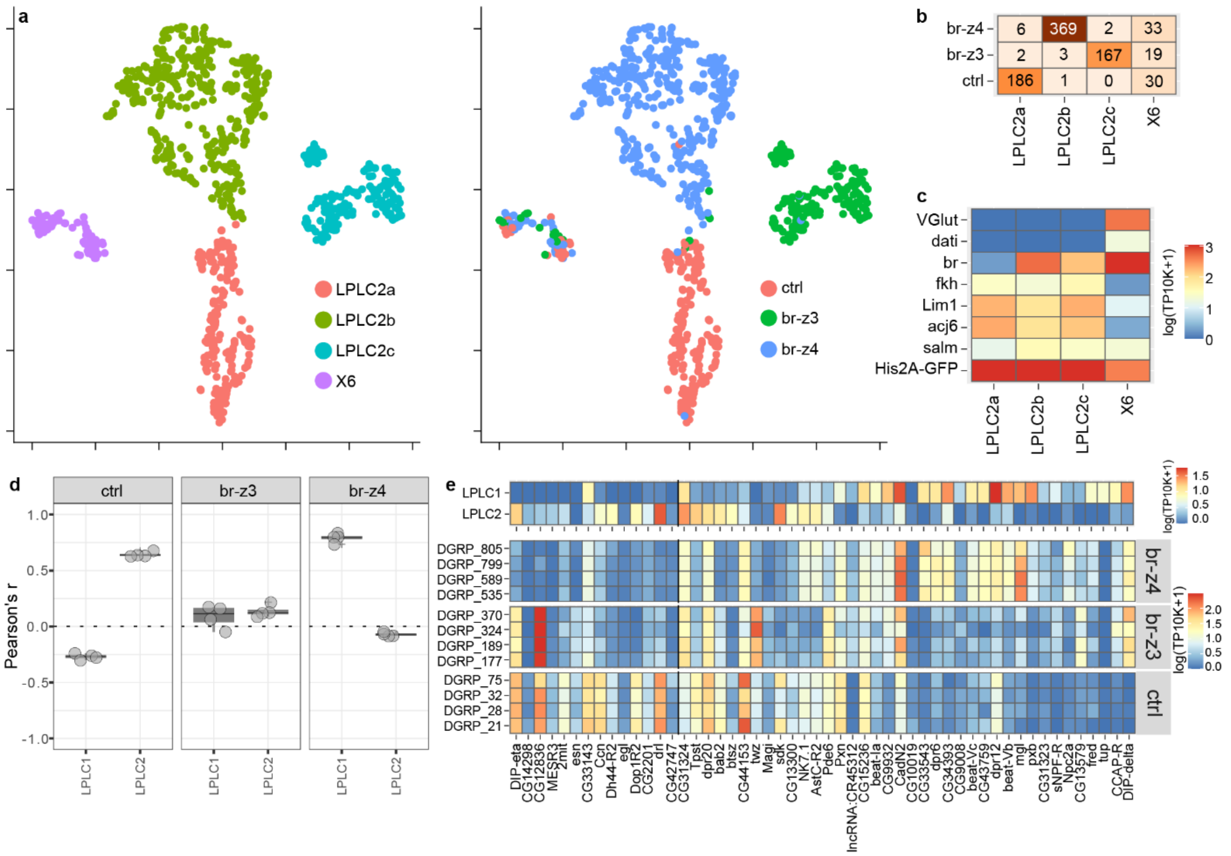


**Supplementary Figure 4: Multiplexed Perturb-seq at 72h APF.**

**(a)** t-distributed stochastic neighbor embedding (tSNE) plots are used for visualization of the clustering of the data. Left, cells are color coded based on unsupervised clustering; Right, cells are color coded based on experimental conditions.

**(b)** Cell counts across clusters and conditions.

**(c)** Expression levels of LPLC2-specific transcription factors (Kurmangaliyev et al., 2020; Ozel et al., 2022) nuclear GFP reporter, and two genes enriched in ectopic, non-LPLC2, clusters (X4/X5).

**(d)** Correlation analysis between transcriptional profiles of LPLC1 and LPLC2 neurons from the developmental atlas (Kurmangaliyev et al., 2020) and LPLC2 neurons in each experimental condition. Comparisons are based on differentially expressed genes (DEGs) between LPLC1 and LPLC2 from **(f)**. Circles are Pearson's r for individual replicates; boxplots are distributions.

**(e)** Heatmaps of expression patterns of DEGs between LPLC1 and LPLC2 at 48h APF. Expression patterns are shown in the atlas (top) and for each condition and replicate. DGRP lines for each X chromosome and replicate are indicated. See Methods for thresholds.

DEGs in **(d)** and **(e)** were identified at 48h APF as there were too few LPLC1/LPLC2 cells in the atlas at 72h APF.
