## Supplemental Tables for "Neuronal identity control at the resolution of a single transcription factor isoform"

Fly Stocks Table

| Name | Designation | Source or Reference | Identifiers |
|---|---|---|---|
| **‘LPLC2-split-GAL4′:** *R19G02_p65ADZp (attP40); R75G12_ZpGdbd (attP2)* | OL0048B | (Wu et al., 2016) | BDSC_48860 |
| **‘LPLC1_1-split-GAL4’:** *R64G09_P65ADZp (attP40); R37H05_ZpGbd (attP2)* | OL0029B | (Wu et al., 2016) | BDSC_47892 |
| **‘LPLC1_2-split-GAL4’:** *R64G09_P65ADZp (attP40); VT045990_ZpGbd (attP2)* | SS02569 | (McFarland et al., 2024) | Robot ID: 3018575 |
| **‘LPLC1_3-split-GAL4’:** *R64G09_P65ADZp (attP40); VT063739_ZpGbd (attP2)* | SS02570 | (McFarland et al., 2024) | Robot ID: 3018576 |
| **smGFP reporter:** *pJFRC200-10XUAS-IVS-myr::smGFP-HA (attP18), pJFRC216-13XLexAop2-IVS-myr::smGFP-V5 su(Hw)attP8* | smGFP | (Nern et al., 2015) | Robot ID: 1116963 |
| **UAS-EGFP:** *P{w[+mC]=UAS-EGFP}8, w[1118]* | UAS-EGFP | Donor: Eric Spana, Duke University; Donor's Source: Martin Zeidler, Harvard Medical School | RRID: BDSC_5428 |
| **LexAop-CsChrimson:** *13xLexAop-CsChrimson-mVenus (attP18)* | LexAop-CsChrimson | Vivek Jayaraman & (Klapoetke et al., 2014) | Robot ID: 1150410 |
| **T2 driver:** *w[1118]; P{y[+t7.7] w[+mC]=GMR47E02-lexA}attP40* | LexA T2 | (Pfeiffer et al., 2010) | RRID: BDSC_53477 |
| **br RNAi-1:** *y[1] v[1]; P{y[+t7.7] v[+t1.8]=TRiP.JF02585}attP2* | br RNAi-1 | (Perkins et al., 2015) | RRID:BDSC_27272 |
| **br RNAi-2:** *y[1] v[1]; P{y[+t7.7] v[+t1.8]=TRiP.HMS00042}attP2/TM3, Sb[1]* | br RNAi-2 | (Perkins et al., 2015) | RRID:BDSC_33641 |
| **UAS br-z1:** *P{w[+mC]=UAS-br.Z1}11-1, w[1118]* | UAS br-z1 | (Zhou et al., 2004) | RRID:BDSC_51379 |

| | | | |
|---|---|---|---|
| **UAS br-z2:** *w[1118]; P{w[+mC]=UAS-br.Z2}20-1* | UAS br-z2 | (Zhou et al., 2004) | RRID:BDSC_51380 |
| **UAS br-z3:** *w[1118]; P{w[+mC]=UAS-br.Z3}216-1* | UAS br-z3 | (Zhou et al., 2004) | RRID:BDSC_51192 |
| **UAS br-z4:** *w[1118]; P{w[+mC]=UAS-br.Z4}37-6* | UAS br-z4 | (Zhou et al., 2004) | RRID:BDSC_51193 |
| **UAS-mCherry:** *y[1] sc[*] v[1] sev[21]; P{y[+t7.7] v[+t1.8]=VALIUM20-mCherry.RNAi}attP2* | UAS mCherry | (Perkins et al., 2015) | RRID:BDSC_35785 |
| **Wild-type background** | CSMH | Martin Heisenberg, University of Wurzburg | |
| **hsFlp:** *w[1118] P{ y[+t7.7] w[+mC]=hs-FLPG5.PEST}(attP3)* | hsFlp construct for MCFO | (Nern et al., 2015) | RRID:BDSC_62118 |
| **FRT:** *w[1118]; P{y[+t7.7] w[+mC]=10xUAS(FRT.stop)myr::smGdP-V5-THS-10xUAS(FRT.stop)myr::smGdP-FLAG}su(Hw)attP5* | FRT construct for MCFO | (Nern et al., 2015) | RRID:BDSC_62124 |
| **DGRP reference panel lines used:**<br>21, 28, 32, 40, 75, 177, 189, 324, 370, 530, 535, 589, 799, 805, 837 | DGRP reference flies | (Huang et al., 2014) | RRID:BDSC_28122, RRID:BDSC_28124, RRID:BDSC_55015, RRID:BDSC_29651, RRID:BDSC_28132, RRID:BDSC_28150, RRID:BDSC_28152, RRID:BDSC_25182, RRID:BDSC_28182, RRID:BDSC_28208, RRID:BDSC_28213, RRID:BDSC_28237, RRID:BDSC_28246 |
| **Nuclear reporter for scRNA-seq:**<br>UAS-H2A-GFP | UAS-H2A::GFP | (Kurmangaliyev et al., 2020) | PMID: 26687360 |
